# P53 and BCL-2 family proteins PUMA and NOXA define competitive fitness in Pluripotent Cells

**DOI:** 10.1101/2023.05.21.541667

**Authors:** Jose A. Valverde-Lopez, Lin Li-Bao, Covadonga Díaz-Díaz, Rocío Sierra, Elisa Santos, Giovanna Giovinazzo, Miguel Torres

## Abstract

Cell Competition is a process by which neighboring cells compare their fitness. As a result, viable but suboptimal cells are selectively eliminated in the presence of fitter cells. In the early mammalian embryo, epiblast pluripotent cells undergo extensive Cell Competition, which prevents suboptimal cells from contributing to the newly forming organism. While competitive ability is regulated by MYC in the epiblast, the mechanisms that contribute to competitive fitness in this context are largely unknown. Here, we report that P53 and its pro-apoptotic targets PUMA and NOXA regulate apoptosis susceptibility and competitive fitness in pluripotent cells. PUMA is widely expressed specifically in pluripotent cells *in vitro* and *in vivo*. We show that the p53-PUMA/NOXA pathway regulates mitochondrial membrane potential and oxidative status. We found that P53 regulates MYC levels in pluripotent cells, which connects these two Cell competition pathways, however, MYC and PUMA/NOXA levels are independently regulated by P53. We propose a model that integrates a bifurcated P53 pathway regulating both MYC and PUMA/NOXA levels and determines competitive fitness through regulation of mitochondrial activity.

## INTRODUCTION

Cell Competition (CC) is a process based on the interaction of neighboring cells of the same type. By this mechanism, less fit cells “*loser cells”* are non-autonomously eliminated upon confrontation with fitter cells called “*winners*”. Cell competition selectively detects and eliminates viable but suboptimal, mis-specified or mis-placed cells, being envisioned as a conserved and extended quality control system in metazoans. From embryonic development to the adult, Cell Competition functions to ensure the proper performance of tissues and organs. In addition, it plays an important role in aging, tissue regeneration and cancer (Clavería & Torres, 2015; Bowling *et al*, 2019; Gregorio *et al*, 2016).

In the early mammalian embryo, the epiblast contains the pool of pluripotent cells that are destined to generate the whole new individual. During pluripotency establishment and through its transition toward differentiation, cells undergo significant alterations in proliferation rate as well as in metabolic, epigenetic and signaling rewiring. Starting at epiblast specification, pluripotent cells are susceptible to undergo apoptosis, with a wave of cell death that peaks at the pre-gastrulation epiblast in a programmed and systematic manner. At this stage, cells become hypersensitive to apoptotic stimuli (Coucouvanis & Martin, 1995; Manova *et al*, 1998; Heyer *et al*, 2000; Pernaute *et al*, 2014, 2022). This apoptosis wave ends with gastrulation, which also coincides with termination of pluripotency. At least in part, this cell death wave results from endogenous Cell Competition that eliminates prematurely differentiating, suboptimal or potentially harmful cells in presence of fitter cells, optimizing the cell pool that will give rise to the new individual (Clavería *et al*, 2013; Sancho *et al*, 2013; Díaz-Díaz *et al*, 2017; Lima *et al*, 2020; Singla *et al*, 2020). In this endogenous CC model, winner cells correlate with high expression of MYC and low expression of p53 transcription factors. Cells with low MYC levels are eliminated by the presence of cells with higher levels, as a mechanism to select metabolically competent cells and removing those cells more prone to differentiate prematurely, safeguarding pluripotency (Clavería *et al*, 2013; Sancho *et al*, 2013; Díaz-Díaz *et al*, 2017). Similar observations apply to the *in vitro* counterparts of epiblast cells as they transit from naïve to primed pluripotency and differentiation. Although, different pathways have been reported to regulate fitness in Pluripotent CC (Sancho *et al*, 2013; Bowling *et al*, 2018; Lima *et al*, 2021; Clavería *et al*, 2013; Hashimoto & Sasaki, 2019; Montero *et al*, 2022), many aspects of the fundamental mechanisms regarding this process remains unknown. In particular, how competitive fitness is encoded in pluripotent cells is not understood.

Identifying pre-existing conditions in prospective loser cells contributing to their loser status would provide insight into the early steps of Cell Competition and fitness comparison. Here, we have explored different factors and pathways regulating cell fitness and also the execution of loser cell death during Pluripotent Cell Competition. As a result, we have identified several candidates of the P53 pathway and propose a model based on increased susceptibility to apoptosis, autophagy and reduction of mitochondrial function, accounting, at least in part, for the loser fitness “*signature”* in Pluripotent Cell Competition. P53 has been identified as a pivotal component in loser cells and its pathway is notably upregulated in loser ES cells. We have found that P53 and the pro-apoptotic BCL-2 proteins PUMA and NOXA regulate apoptosis susceptibility in ESCs, in which their function and expression is not restricted to apoptotic cells but present in all the pluripotent cell population. We found that their expression levels correlate with Competitive fitness and their inhibition is enough to promote the winner phenotype in mouse ESCs. P53 regulation of competitive fitness depends on the pluripotency status, with the pathway being activated as the cells progress towards differentiation, which increases apoptotic hypersensitivity and their ability to induce Cell Competition and apoptosis is suppressed in naive pluripotency conditions. Additionally, we have shown that P53 activity inhibits MYC expression and is strictly required for PUMA expression.

We propose a model that integrates the P53 pathway and MYC in the definition of the loser cell fitness “status” and suggests that an alteration in mitochondrial function regulated by BCL2-family proteins underlies competitive fitness in pluripotent cells.

## RESULTS

### P53 pathway is upregulated in MYC-low cells

To identify pathways involved in the regulation of cellular fitness and the execution of loser cells elimination, we compared the transcriptional profile of low- and high-MYC expressing cells. In a previous work, we performed transcriptomic studies comparing low-, medium- and high-MYC expressing cells (described in (Díaz-Díaz *et al*, 2017)). We took advantage of a GFP-MYC reporting cell line, in which GFP levels reliably reports MYC expression sorted cells by FACS according to GFP expression levels and sequenced the transcriptome (Fig. 1A). This procedure allowed us to study candidate genes in the MYC-low cell population, potentially involved in their low competitive ability or responsible for their elimination. We reanalyzed the data in (Díaz-Díaz *et al*, 2017) and used gene-set enrichment analysis to identify P53 as the most enriched pathway in MYC-low cells (Fig. S1A). To validate this correlation between P53 and MYC-low cells, we performed an immunostaining in ES cells. P53 exhibited a heterogeneous nuclear pattern in ES cells (Fig. 1B, C, S1B) and showed a strong increase upon treatment with etoposide (widely P53 activator through DNA damage generation) as a positive control (Fig. S1B). Per-cell quantification of P53 and MYC expression confirmed an inverse correlation between the two proteins (Fig. 1B-D). Then, we checked the apoptotic role of P53 in ES cells by using an anti-active CASP3 antibody. To avoid problems of apoptosis-associated autofluorescence, only cells that maintain an integral cellular morphology (early apoptotic cells) were considered for this quantification (Fig. 1E). Activated CASP3-positive cells displayed higher levels of P53, indicating a correlation between P53 and apoptosis (Fig. 1F). Additionally, when considering CASP3 negative cells only, P53 expression was still higher in MYC-low cells than in MYC-high cells (Fig. 1F) (Díaz-Díaz *et al*, 2017). These observations suggest a role for P53 pathway in the execution of loser cell death but also indicate that P53 could exert a role in defining fitness and the loser “status”. Therefore, we focused on selecting candidate genes from the P53 pathway involved in apoptosis/cell stress, upregulated in MYC-low cells.

**Figure 1.**
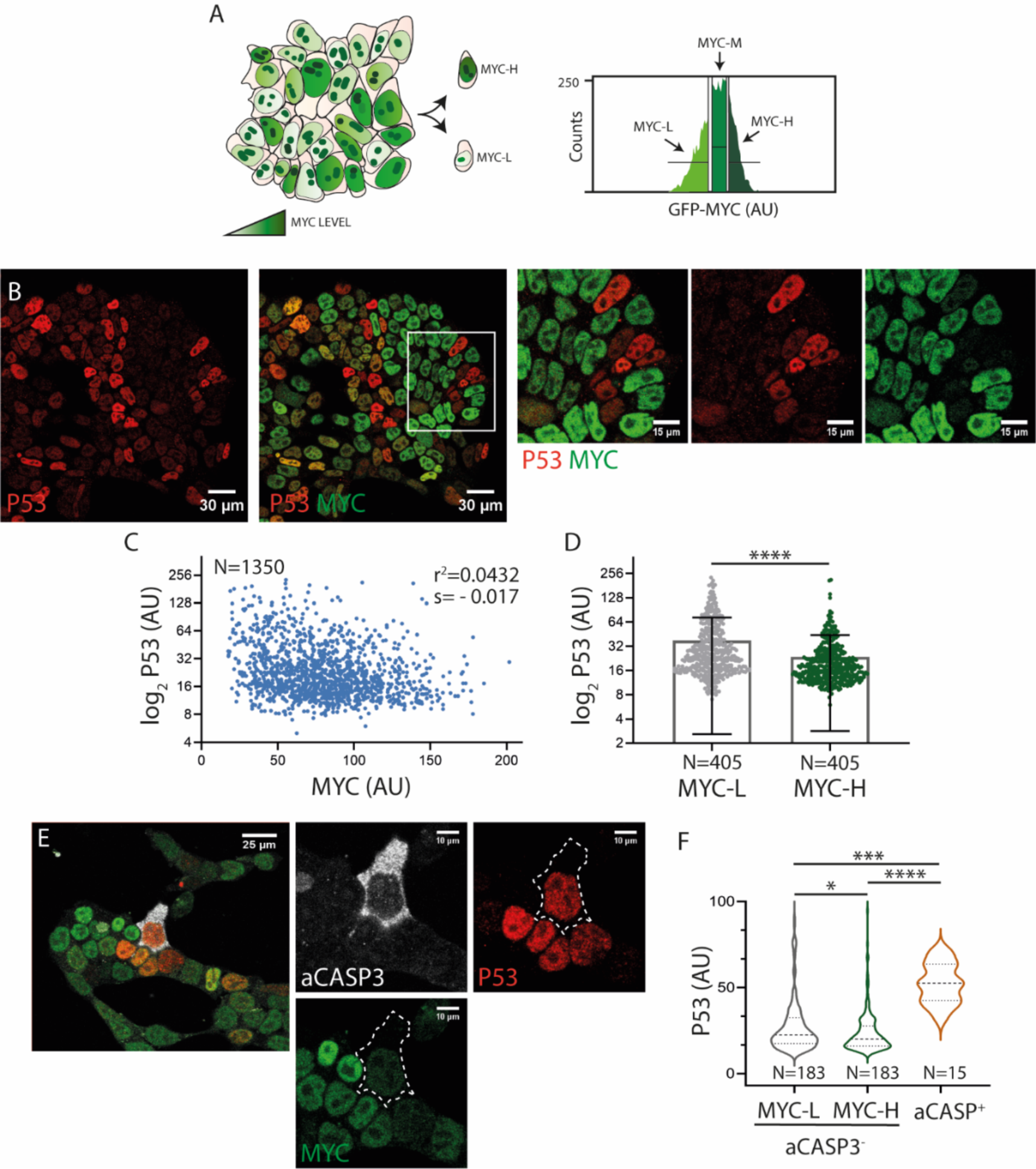
The P53 pathway is upregulated in MYC-low ES cells. **A.** Schematic representation of the GFP-MYC ES cell line (left). Histogram showing the segmentation of GFP-MYC ES cells into three populations (MYC-low, -medium and -high), which were sorted by FACS (right). **B.** Confocal images showing the expression of P53 and MYC in ES cells and magnification (right). **C.** Quantification of P53 and MYC levels, p<0.0001. **D.** P53 levels in MYC-low and MYC-high populations. **E.** Confocal images showing active CASP3, P53 and MYC immunostaining and magnification (right). **F.** Quantification of P53 in active CASP3 positive cells and in MYC-low and MYC-high aCASP3 negative cells.

By using gene ontology (GO) terms related to P53 pathway and apoptosis we were able to select those genes involved in the P53 pathway and apoptosis from our RNAseq data (GO terms are described in Materials and Methods). We identified some genes such as *trp53inp1*, *ddit4* or *perp*. Moreover, we found several members of the apoptotic family protein BCL-2 (Fig2A, Fig S1C). The overexpression of these candidate genes in MYC-Low cells was analyzed by qPCR obtaining similar results (Fig. S1D).

*Trp53inp1* is a stress-induced protein that induces autophagy and mitophagy (Saadi *et al*, 2015). *Ddit4* is upregulated upon stress and affects mitochondrial biogenesis and metabolism. It functions as a strong inhibitor of mTORC1, which induces autophagy (Tirado-Hurtado *et al*, 2018). *Perp* encodes a plasma membrane protein which can interact with the Ca^2+^ pump (SERCA2B) in the endoplasmic reticulum, inducing apoptosis (McDonnell *et al*, 2019). Finally, we identified different genes belonging to the BCL-2 family protein (Fig2B).

BCL-2 (B cell lymphoma-2) proteins constitute important regulators of apoptosis. Structurally, these proteins possess a conserved BH domain (BCL-2 homology), critical for their function and they are classified as multi-BH proteins, including anti-apoptotic proteins (BLC-2, BCL-XL, MCL-1) and pro-apoptotic proteins (BAX, BAK, BOK) or with a single BH domain, “BH3-only proteins”, which exert a pro-apoptotic role (BIM, BAD, tBID, NOXA, PUMA) (Fig. S2A). Upon apoptotic stimuli, multi-BH pro-apoptotic proteins BAX, BAK and BOK can oligomerize and generate pores in the mitochondrial outer membrane (MOM) allowing pro-apoptotic factors to release and trigger the apoptosis. This oligomerization is tightly controlled by the balance between anti- and pro-apoptotic BCL2 proteins (Certo *et al*, 2006) (Fig. S2B).

From this family, we analyzed PUMA (*bbc3*) expression, one of the most important apoptotic factors downstream P53 (Yu & Zhang, 2008). PUMA was expressed in almost all ESCs by immunostaining, exhibiting a cytosolic pattern with variable levels of expression (Fig. 2C). Per-cell quantification of PUMA and MYC levels revealed an inverse correlation, which was confirmed by immunoblot (Fig. 2D, E & Fig. S2C). Then, we performed an aCASP3 staining, showing that apoptotic cells expressed moderately higher PUMA levels with respect to the general cell population (Fig. 2F, G). We found that *puma* upregulation in MYC-low cells corresponds to the main isoform, isoform 1 (Fig. S2D).

**Figure 2.**
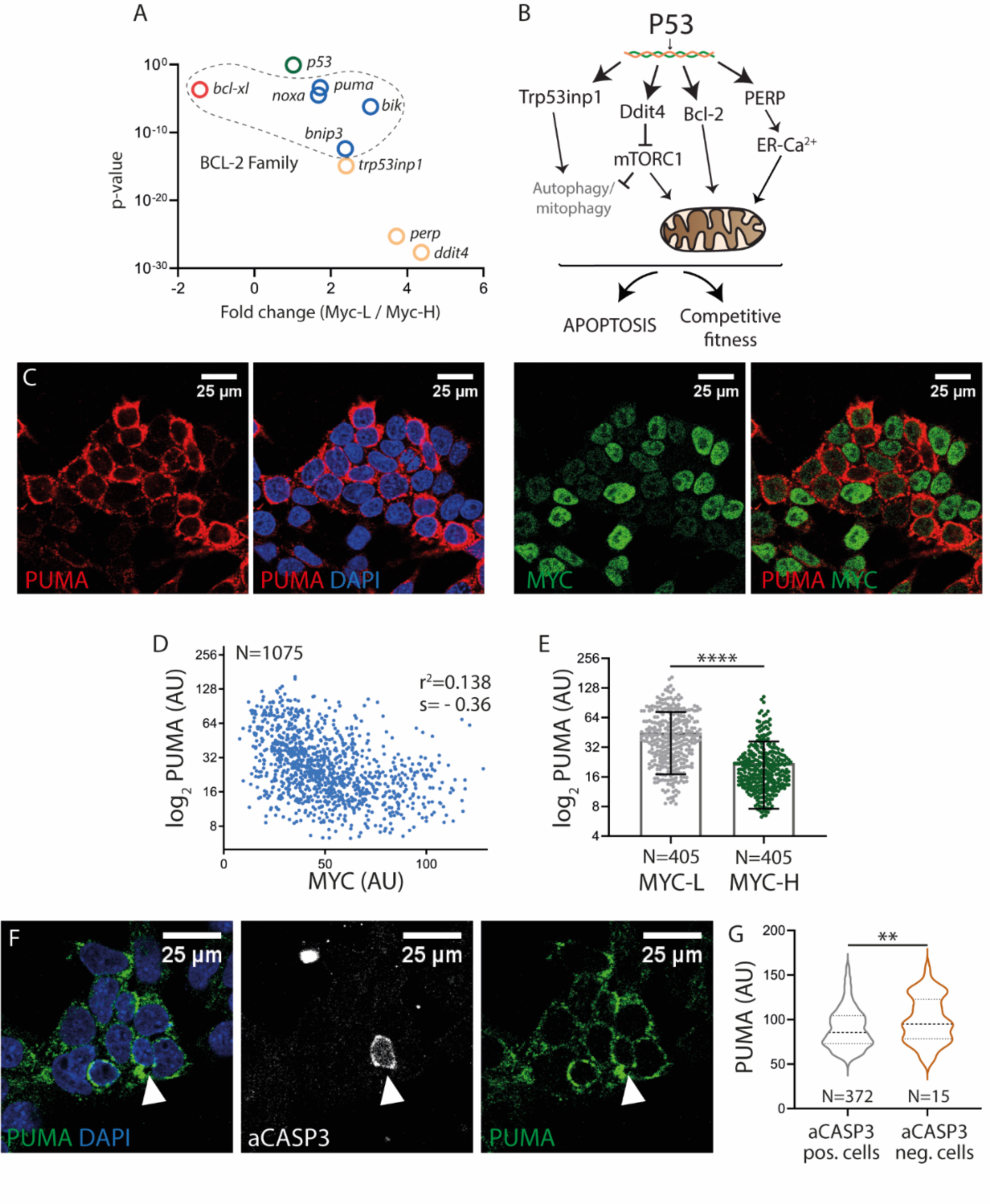
Candidate genes upregulated in MYC-low cells. PUMA levels inversely correlate with MYC levels. **A.** Dot plot showing the fold change and p value of candidate genes. **B.** Schematic representation of different candidate genes related to the P53 pathway and involved in apoptosis/stress, upregulated specifically in MYC-low cells. **C.** Confocal images showing PUMA and MYC expression in ESCs. **D, E**. Quantification of PUMA and MYC expression, p<0.0001 and expression of PUMA in MYC-L and MYC-H cells. **F.** Confocal images of aCASP3 and PUMA immunostaining. **G.** Quantification of PUMA levels in aCASP3 positive and negative cells.

The fact that PUMA is heterogeneously expressed in almost all cells and that apoptotic cells showed just a moderately increased in PUMA levels suggest that although PUMA and P53 have a role regulating apoptosis, PUMA may exert another function in ES cells. Considering the inverse correlation between PUMA and MYC, and that MYC is a well-described fitness regulator, we hypothesize that PUMA could play a role in regulating competitive fitness.

### P53-PUMA and MYC regulation

First, we studied the regulatory interactions between P53 and PUMA. To do so, we performed an immunostaining against P53, PUMA and MYC proteins and we established a positive correlation between P53 and PUMA in ESCs (Fig. 3A, B), both of which inversely correlate with MYC levels (Fig. 3A, C). The activation of P53 using etoposide efficiently upregulated PUMA levels (Fig. S3A, B). Then, we generated a *p53* knockout ES cell line and checked PUMA expression. Notably, in the absence of P53, we observed no detectable PUMA signal in ES cells (Fig. 3D), indicating that P53 is required for PUMA expression.

**Figure 3.**
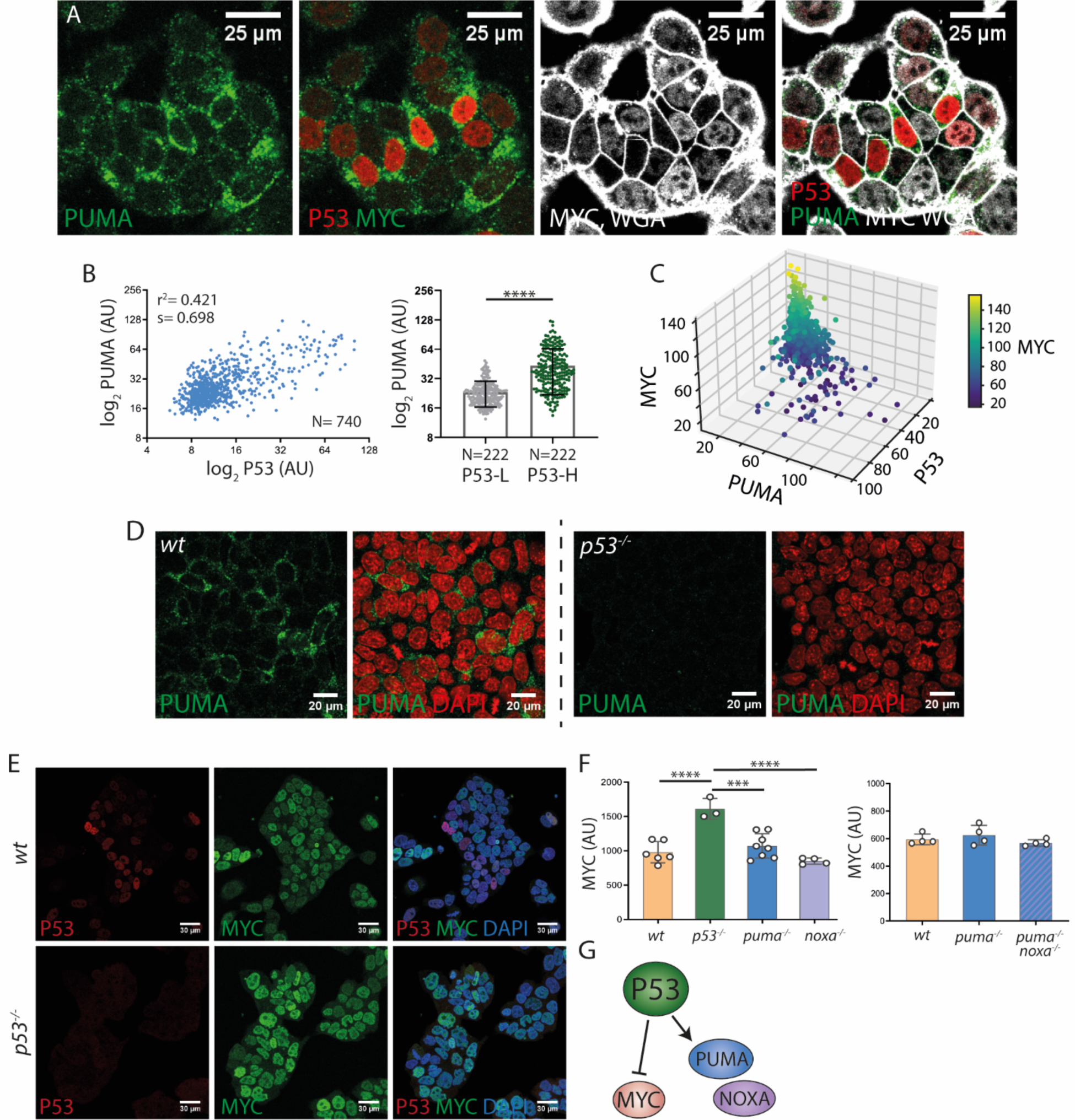
P53 regulation of PUMA and MYC in ES cells. **A.** Confocal images showing P53, PUMA and MYC expression. **B.** Quantification of P53 and PUMA expression. **C.** Quantification of P53, PUMA and MYC levels, N=384, PUMA-MYC (r^2^=-0.506, p=2.09×10^-26^), P53-MYC (r^2^=-0.457, p=2.09×10^-21^), P53-PUMA (r^2^=0.669, p=3.50×10^-51^). Graph generated with Python. Heatmap corresponds to MYC expression. **D.** Confocal captures showing PUMA levels in *wt* and *p53^-/-^* cells. **E.** Confocal images of P53 and MYC levels in *wt* and *p53^-/-^* cells. **F.** Quantification of MYC levels in *wt, p53^-/-^, puma^-/-^, noxa^-/-^* (left) and *wt*, *puma^-/-^*and the DKO *puma^-/-^*and *noxa^-/-^*(right). **G.** Schematic representation of P53 regulating the expression of PUMA and NOXA while exerting an inhibitory effect over MYC expression.

Next, we wanted to explore the regulatory interactions between MYC and the P53-PUMA pathway. First, we analyzed the levels of P53 and PUMA using a *Myc* knockout cell line and we found that *Myc* deletion did not increase P53 or PUMA expression, but rather we observed a slight non-significant downregulation (Fig. S3C, D). Subsequently, we examined the expression of P53 and PUMA when MYC is overexpressed, taking advantage of the mouse ES cells carrying the *iMOS^MYC^* allele (Clavería *et al*, 2013). This cell line allows us to induce with tamoxifen a population of EYFP positive cells overexpressing MYC and a population of wild type ECFP cells in a random mosaic manner (Fig. S3E). We found that MYC overexpression did not decrease the levels of P53 or PUMA (Fig. S3E). These results show that MYC does not regulate P53-PUMA expression.

Then, we analyzed whether, alternatively, P53 regulates MYC. We found that in P53-deficient cells, MYC was upregulated (Fig. 3E, F). Additionally, activation of P53 using Nutlin3 (an extensively characterized P53 activator) led to MYC downregulation (Fig. S4A-C). In similarity to etoposide, Nutlin3 led to the upregulation of PUMA expression.

Next, we studied whether PUMA (as well as other BH3-only proteins such as NOXA (*pmaip1*)), also play a role in MYC regulation. In the absence of either PUMA or NOXA, or both we did not find a significant change in MYC expression (Fig. 3E, F). These results indicate that P53 acts upstream both PUMA and MYC. P53 is required for PUMA expression and P53 inhibits MYC expression, but not through the regulation of BH3-only proteins PUMA or NOXA, (Fig. 3G).

We then wanted to explore P53-PUMA and MYC correlation *in vivo* in the early mouse embryo. E6.5 mouse embryos were used since MYC-driven Cell Competition and other CC models have been described at this developmental stage (Bowling *et al*, 2018; Clavería *et al*, 2013; Díaz-Díaz *et al*, 2017; Sancho *et al*, 2013; Lima *et al*, 2021). Consistent with the observations in ES cells, epiblast cells expressed heterogeneous levels of PUMA. In contrast to the epiblast, the extraembryonic ectoderm (Ex) did not show detectable PUMA expression, while MYC is strongly expressed (Fig. 4A). We found that epiblast cells with high PUMA levels exhibit lower MYC levels than the general cell population (Fig. 4B, C). Additionally, we found no detectable PUMA signal in *p53^-/-^* embryos (Fig. 4D) indicating that P53 is essential for PUMA expression in the mouse embryo.

**Figure 4.**
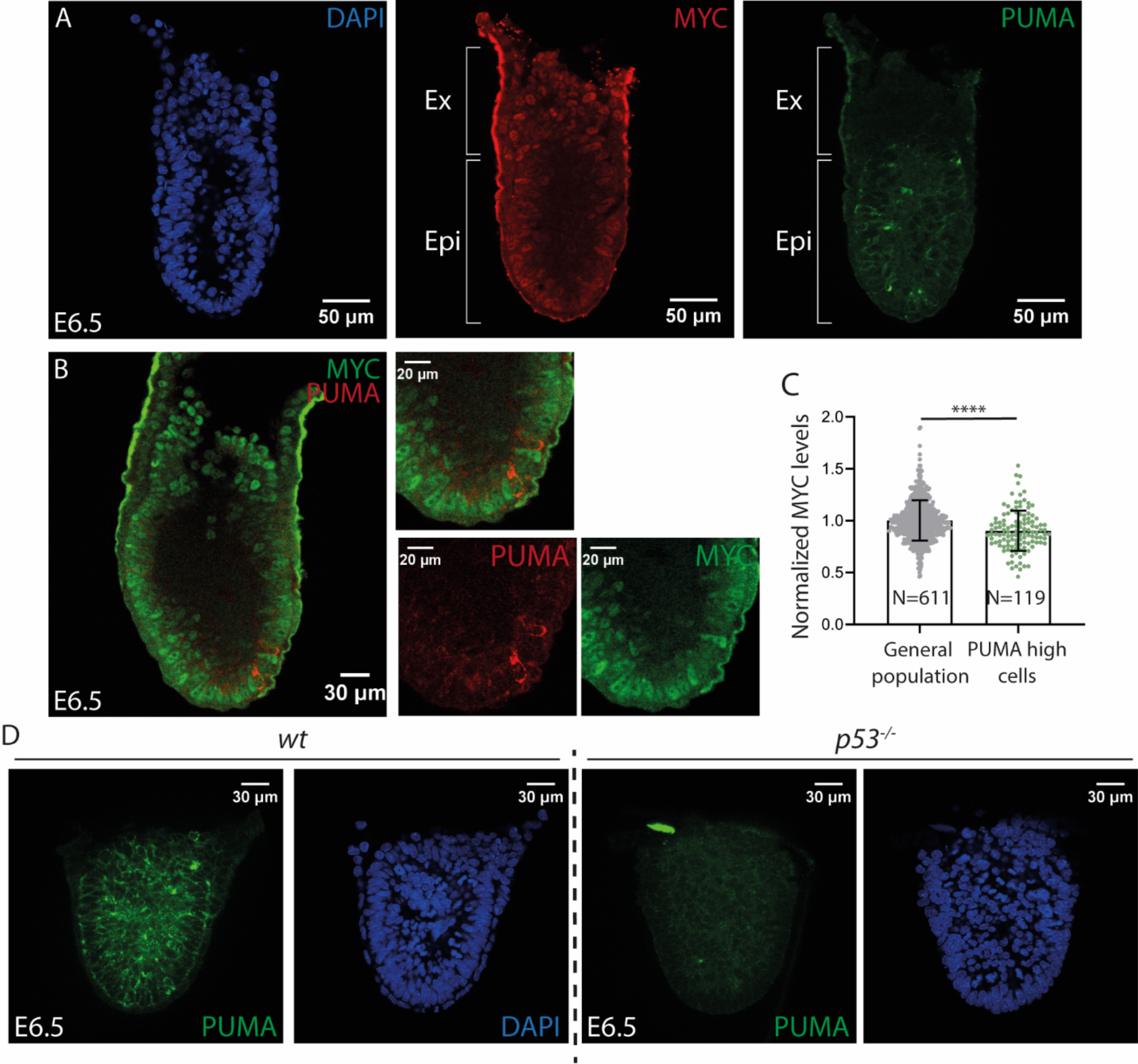
P53, PUMA and MYC expression and regulation in the early mouse embryo. **A, B.** Confocal images showing PUMA and MYC expression in an E6.5 mouse embryo (Ex, extraembryonic ectoderm; Epi, epiblast). **B**. Quantification of normalized MYC levels in the general cell population and high PUMA-expressing cells. **C.** Confocal images showing PUMA expression in *wt and p53^-/-^* E6.5 embryos.

Subsequently, we wanted to explore P53 expression in the epiblast and its correlation with PUMA and MYC. Although P53 is present and functional at E6.5 (since PUMA expression is not detected after *p53* deletion), we detected almost no P53 positive cells, even upon activation with Nutlin-3. Therefore, we turned to a previous stage, E3.5, where CC interactions have also been described (Hashimoto & Sasaki, 2019). At this stage, P53 showed a nuclear pattern analogous to ES cells. We found a positive correlation between P53 and PUMA (Fig S5A, B), but no correlation was found between P53 and MYC (Fig S5C, D). Eventually, we explored whether P53 exerts an inhibitory effect over MYC. However, we did not observe an upregulation of MYC in neither *p53*^-/-^ E6.5 embryos (Fig S5E) nor E3.5 embryos (Fig S5F). These results suggest that in the early embryo, MYC regulation by P53 is not as simple as in cultured ESCs and additional factors may add complexity to the regulatory network.

### P53-PUMA and MYC are regulated accordingly to the pluripotency status

We previously described that MYC is regulated by the pluripotency status, thus, we explored the relationship between Pluripotency status and the P53-PUMA pathway. Pluripotency is defined as the capacity of embryonic cells to self-renew and generate all embryonic lineages. Although pluripotent cells maintain a core pluripotency TF regulatory network, pluripotency is not a single status, but a set of dynamics stages in which cells change their gene expression, epigenetic landscape and metabolic profile in a continuous manner during development (Nichols & Smith, 2009; Sperber *et al*, 2015). In the mouse embryo, pluripotency comprises from E3.5 to E6.5-E7.5. At E3.5, the pluripotent cells present a “naïve” pluripotent status characterized by a generalized hypomethylated “open” chromatin. Cells then evolve towards a “primed” pluripotent status through a process called “formative pluripotency” (Kinoshita *et al*, 2021). Primed pluripotent cells have gained extensive methylation marks and have already established the X-chromosome inactivation. Eventually, pluripotency ends as cells differentiate at E6.5-E7.5 into the three germ layers during gastrulation (Hackett & Surani, 2014; Posfai *et al*, 2014) (Fig S6A).

Different pluripotent states can be recreated *in vitro*. The use of two differentiation inhibitors, PD03 and CHIRON maintains cells in a “naïve” status (the so-called “2i medium”) (Ying *et al*, 2008; Sato *et al*, 2004). The use of Activin A and FGF transiently promotes the formative status and eventually its transition to the primed state (Fig S6A) (Tesar *et al*, 2007; Brons *et al*, 2007), while the use of serum + LIF or SR + LIF (KO Serum Replacement, which is a chemically defined formula that substitutes serum) an promotes a mix of naïve and primed cells (Guo *et al*, 2016).

Therefore, we analyzed P53 and PUMA expression dynamics in different pluripotency culture conditions: 2i medium, which promotes a “naïve” status, conventional medium (Fetal Bovine Serum (FBS) + LIF) and Serum Replacement medium (SR+LIF). We found that P53 and PUMA levels increased expression and per-cell variability as cells transit from a naïve into mixed pluripotency. A similar regulation affects MYC expression (Fig. 5A, C). Furthermore, allowing differentiation by removing LIF, led to a decrease in MYC levels, while PUMA increased (Figure S5B). Collectively, these results indicated that P53 and PUMA are regulated by pluripotent conditions and their levels increase as cells progress towards the formative pluripotency.

**Figure 5.**
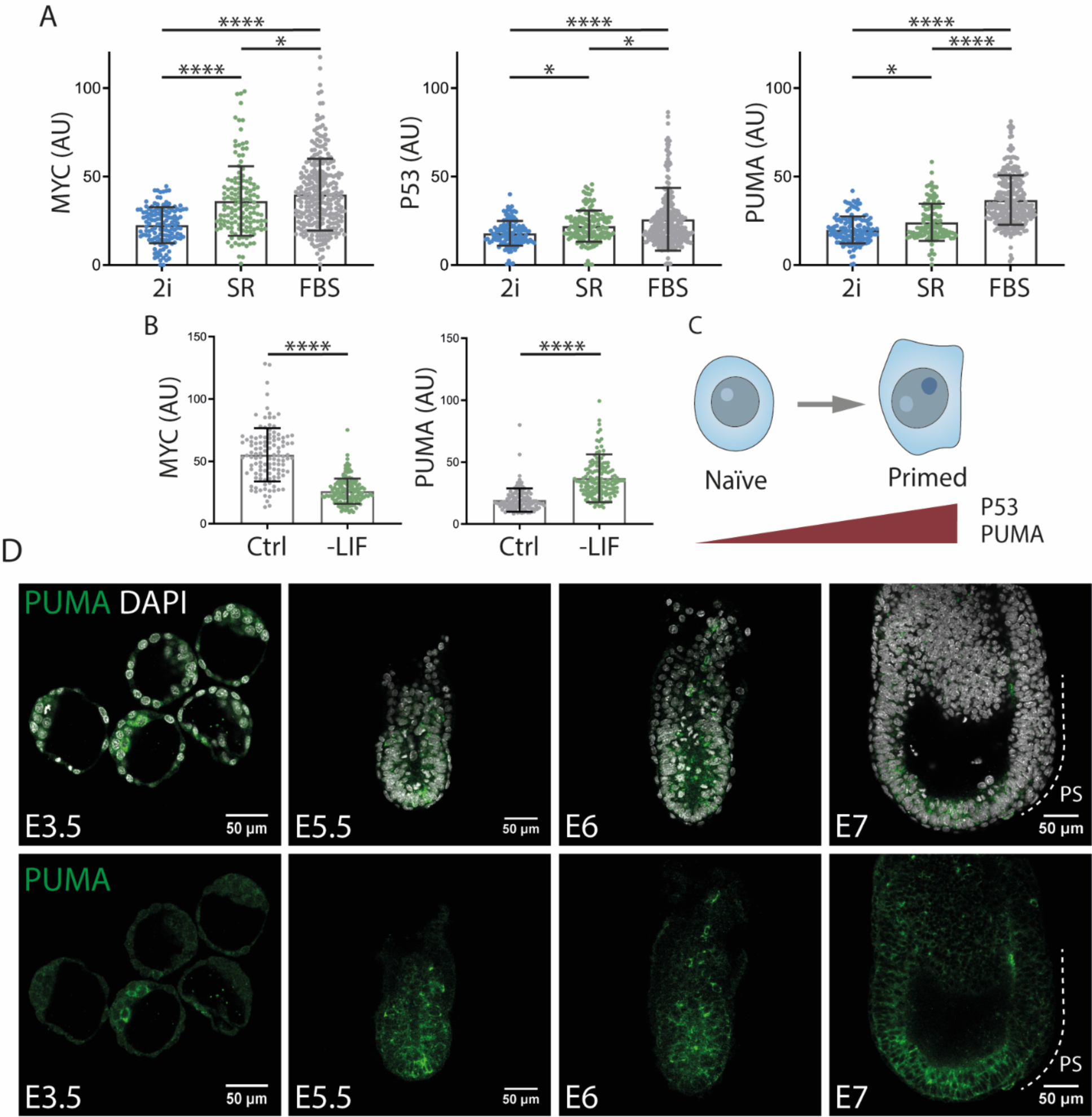
The pluripotency status regulates P53 and PUMA expression. **A.** Quantification of MYC, P53 and PUMA in 2i, SR and FBS conditions. **B.** Quantification of MYC and PUMA expression in conventional or differentiating conditions (removing LIF). **C.** Scheme of the evolution of P53 and PUMA expression as ES cells evolve from naïve to formative status. **D.** Confocal images displaying PUMA expression pattern in mouse early embryos at the signaled stages.

Next, we studied the expression pattern of PUMA during early mouse embryo development. PUMA was already detected E3.5 at heterogeneous levels, with some ICM cells showing high expression levels. Subsequently, from E5 to E6.5, PUMA is heterogeneously expressed in the epiblast, in similarity to ES cell cultures. Then, when gastrulation begins, PUMA levels strongly decreased in the gastrulating cells of the primitive streak (Fig. 5D). A pattern of high expression of PUMA in a heterogeneous pattern therefore seems to be related to pluripotency *in vitro* and *in vivo*.

### Functional characterization of P53 and the BH3-only proteins PUMA and NOXA

After analyzing the P53-PUMA and MYC regulation, we studied their function in ES cells. P53 is well known for inducing cell cycle arrest and apoptosis in response to DNA damage and more recently it has been also related to other functions such as autophagy, metabolism or differentiation (Kastenhuber & Lowe, 2017). On its side, BCL-2 proteins are mostly related to apoptosis, although other functions such as metabolic regulation have been recently described (Siddiqui *et al*, 2015; Kim *et al*, 2019). In ES cells, the role of P53 in apoptosis and cell cycle arrest is not clear and recent works suggest that P53 functions vary as ES cells evolve through the pluripotent states (Fu *et al*, 2020; Hao *et al*, 2020; Jaiswal *et al*, 2020). Regarding PUMA and NOXA, their role in Pluripotent Stem Cells is for the most part unknown. To test the role of P53, PUMA and NOXA in apoptosis, cell cycle arrest and differentiation, we used knockout ESC lines. By performing an immunostaining against aCASP3, we found that in the absence of either P53, PUMA or NOXA there was a decrease in apoptosis (Fig 6A, B). Similar results were obtained by using a fluorogenic substrate of CASP3/7 (FLICA^TM^) (Fig S7A).

**Figure 6.**
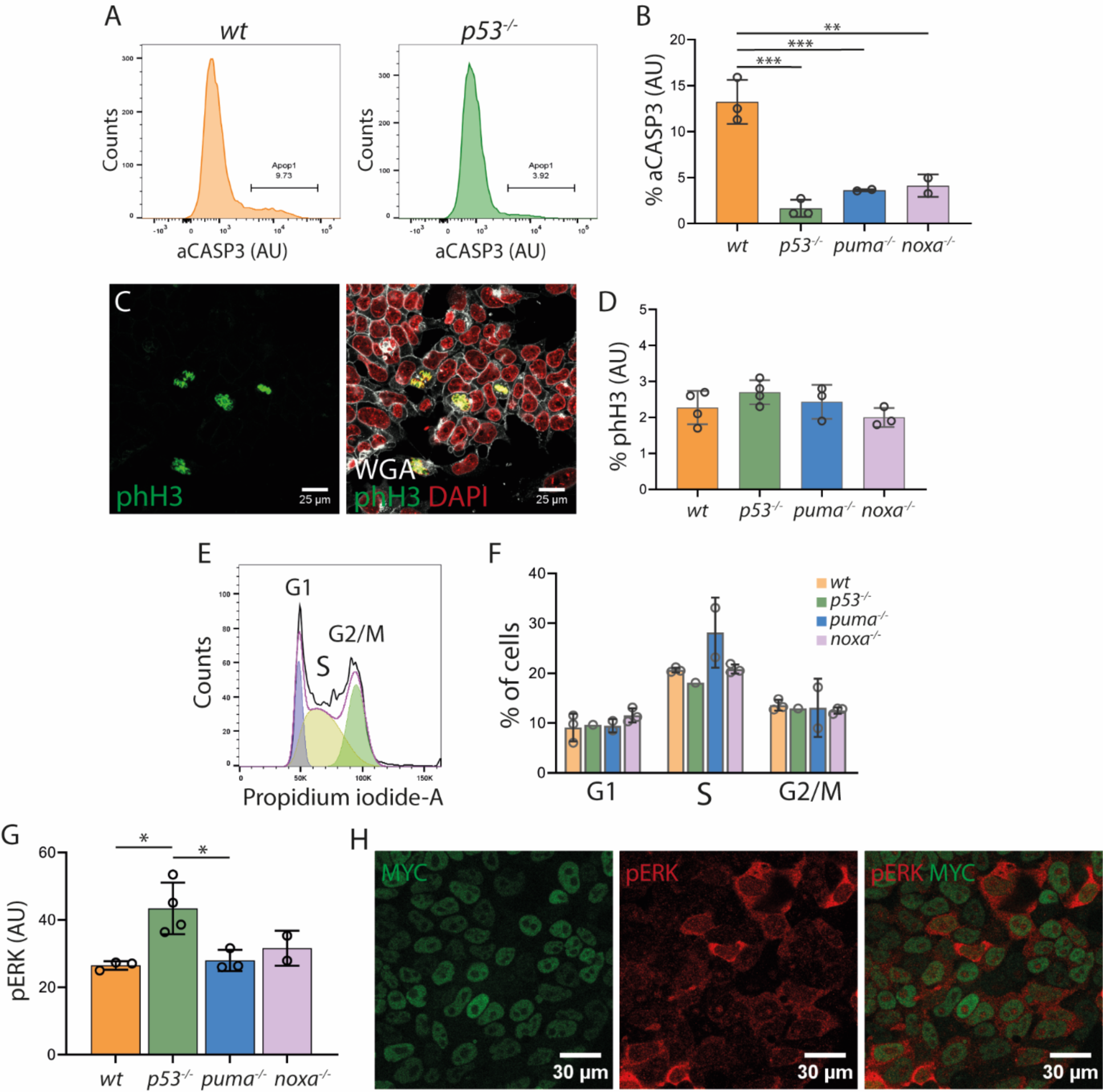
Role of P53 and BH3-only protein PUMA and NOXA in ESC apoptosis, proliferation and cell cycle. **A.** Histograms showing aCASP3 staining in *wt* and *p53**^-/-^*** cells. **B.** Quantification of the percentage of aCASP3 positive cells. **C.** Confocal images showing phH3 positive cells and quantification (**D**). **E.** Histogram showing cell cycle and quantification (**F**). **G.** Bar graph showing p-ERK levels and confocal captures of p-ERK and MYC expression in ESCs (**H**). “aCasp3” refers to the active or cleaved Caspase3 protein. “phH3” refers to the phosphorylated Histone3 protein.

We then evaluated the proliferation and cell cycle by analyzing phospho-Histone 3 (pH3) immunostaining and propidium iodide staining respectively. The absence of either P53, PUMA or NOXA did not lead to significant changes in proliferation (Fig 6C, D; Fig S7B) or cell cycle (Fig 6E, F).

We finally explored how the mutation of P53 and its targets affects ES cell differentiation status. We found that p-ERK, a marker of primed ESCs (Deathridge *et al*, 2019), is expressed at higher levels in P53-deficient cells than in control ESCs (Fig 6G). In the same test, as previously described (Díaz-Díaz *et al*, 2017), Myc and pERK show an inverse correlation (Fig. 6H). This suggests a role of P53 in ESC transition from pluripotency to differentiation.

### P53 and BH3-only proteins PUMA and NOXA regulate cell competitive fitness

Here, we have identified *p53* and BH3-only members *puma* and *noxa* as genes upregulated in MYC-low cells during Cell Competition and we have reported a role for these factors in apoptosis in ESCs. Considering that P53 and PUMA are expressed in almost every cell and the fact that they exhibit a strong inverse correlation with MYC levels suggests they can contribute to define competitive fitness. As opposed to a pro-apoptotic factor simply involved in the culling of loser cells, a fitness component is expected to change the dynamics of cell competition in a cell non-autonomous manner. To study this aspect, we studied how the elimination of these factors affect the viability of neighboring cells in competition assays. We performed these experiments in cells starting differentiation, as at this stage, the competitive interactions are enhanced (Dejosez, 2013; Lima *et al*, 2020; Sancho *et al*, 2013; Bowling *et al*, 2018).

To do so, we confronted *tdtomato-*expressing *wt* cells with either *p53, puma* or *noxa* knockout cells and with non-fluorescent *wt* cells as a control. During CC assays, we compared the evolution of *tdtomato*-*wt* cells in co-culture with *wt* or the different knockout cell lines (Fig. 7A-C). Additionally, we analysed the evolution of each knockout cell line growing in homotypic conditions (Fig. 7B). *Tdtomato-wt* cells were eliminated when co-cultured together with *p53^-/-^* cells but not when they were co-culture with other *wt* cells (Fig. 7D, left). The population of *tomato*-*wt* cells was also reduced when confronted with *puma^-/-^* cells (Fig. 7E, left). Same experiments with *noxa^-/-^* cells resulted in a non-significant tendency towards a reduced growth of the confronted *tomato*-*wt* cells (Fig. 7F, left). Notably, double knockout *puma^-/-^; noxa^-/-^* cells produced a stronger reduction in the population of *tomato*-*wt* cells than single *noxa^-/-^* or *puma^-/-^* cells (Fig. 7G, left). The differences in the growth of the cell populations can be also observed by the ratio between the final and initial cell number for each population (Figure 7D-G, right). These results indicate that P53 and BH3-only proteins PUMA and NOXA can regulate the Competitive fitness in ES cells in such a way that lower levels of P53, PUMA or NOXA results in higher fitness.

**Figure 7.**
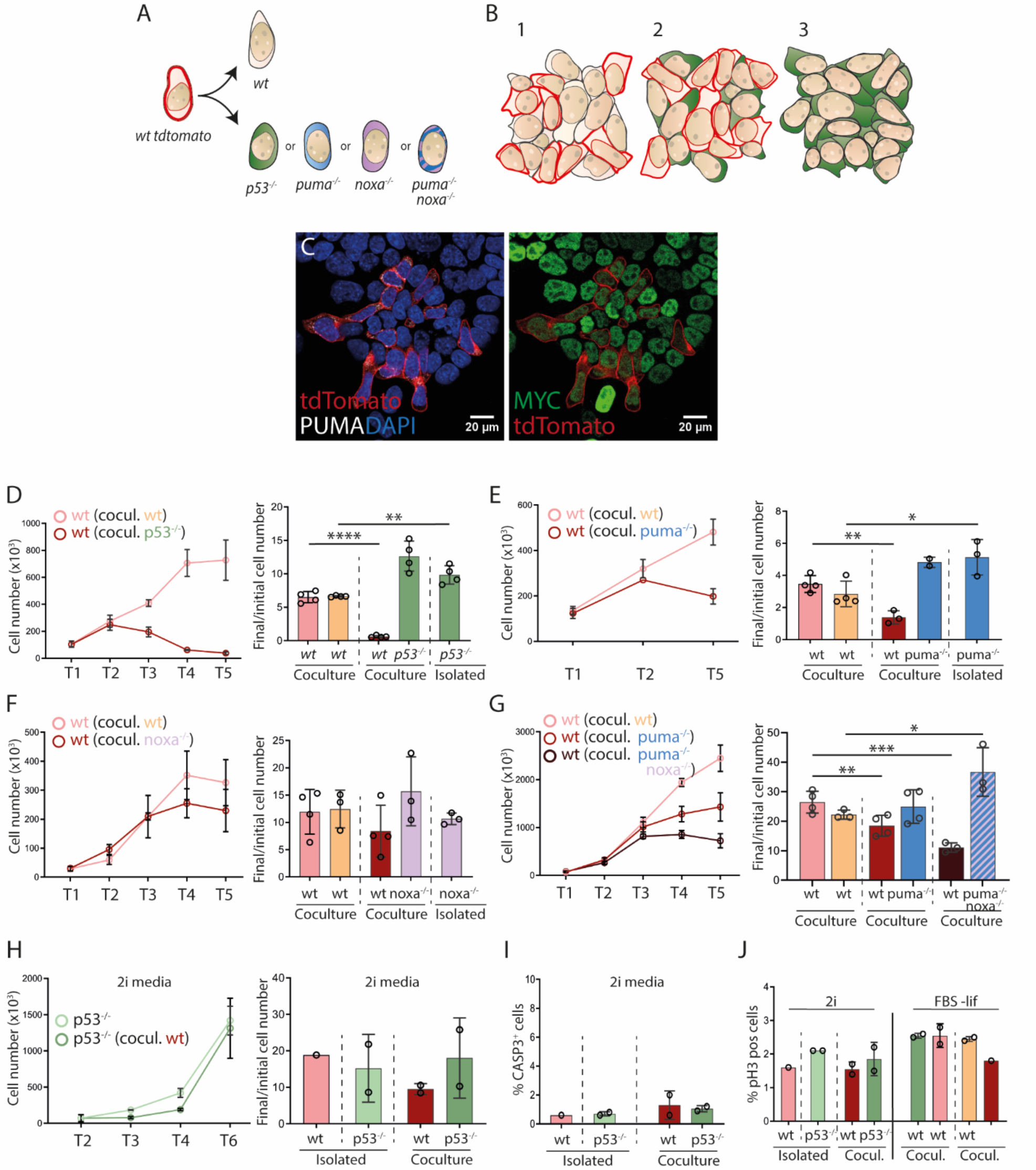
P53, PUMA and NOXA regulate competitive fitness. **A.** Schematic representation of the experimental design. *tdTomato*-*wt* cells were confronted either with knockout cells (represented in different colors) or non-fluorescent *wt* cells as a control. **B.** Schematic representation showing the three types of cell culture we monitor in this assay. (1) td*Tomato*-*wt* in co-culture with non-fluorescent *wt* cells, (2) td*Tomato*-*wt* in co-culture with the different knockout cell lines. Here *p53^-/-^* cells, used as an example of knockout cell line, are represented in green. (3) Homotypic culture of the different knockout cell lines. **C.** Confocal image showing *tdTomato*-*wt* and *puma* knockout cells. **D-G** (left). Evolution of *tdTomato*-*wt* cell number in co-cultured with *wt* cells (light red line) or *p53^-/-^* cells (dark red line) **(D)**, *puma^-/-^***(E)**, noxa*^-/-^* **(F)** or double knockout *puma^-/-^*noxa*^-/-^* cells **(G)**. **D-G** (right). Bar graphs representing the ratio between the final and initial cell number of *wt* and *p53^-/-^, puma^-/-^*, *noxa^-/-^* or the double knockout *puma^-/-^,* noxa*^-/-^* cells in isolated or in co-culture conditions. Each dot represents a different clone. **H** (left). Evolution of *p53^-/-^* cells isolated or in co-culture with *wt* cells. **H** (right). Final/initial cell number ratio of *p53^-/-^* cells and *wt* cells isolated and in co-culture. **I.** Percentage of aCASP3 positive *wt* and *p53^-/-^*cells in the indicated conditions **J.** Percentage of phH3 positive *wt* and *p53^-/-^* in the indicated conditions.

It has been reported in naïve conditions Cell Competition does not take place, so cells start differentiation to undergo competitive interactions (Sancho *et al*, 2013; Bowling *et al*, 2018; Montero *et al*, 2022). In some reports it has even been reported that in naïve conditions *p53^-/-^* cells can switch from winner to a loser behavior (Dejosez, 2013). In agreement with Montero and colleagues (Montero *et al*, 2022), we found that in 2i medium, *p53^-/-^* behave neutrally with respect to *wt* cells; while they do not outcompete *wt* cells, they are neither outcompeted by them (Fig. 7H).

Interestingly, in naïve conditions, *wt* and *p53^-/-^* ESCs show a similar incidence of cell death (Fig. 7I). Additionally, in these conditions, *p53^-/-^* cells do not exhibit the higher growth rate that they showed under pro-differentiating conditions (Fig. 7D and 7H right). Proliferative activity was lower in 2i than in differentiating conditions and again, we did not find differences between WT and *p53^-/-^*cells (Fig. 7J).

These results indicate that P53 in naïve conditions does not regulate cell death or growth rate. This goes in agreement with the low expression of P53 that we have reported in naïve ESCs compared to primed ones (Fig 5A).

### P53 and PUMA cellular functions related to competitive fitness

We then explored the function of P53 and PUMA in fitness regulation and Cell Competition. First, we took advantage of super-resolution STED microscopy to describe PUMA location within the cell and found that it strongly co-localizes with the mitochondria (Fig. 8A). Interestingly, PUMA has been recently reported to promote a metabolic switch regulated by P53 towards glycolysis by inhibiting pyruvate uptake into the mitochondria in human cancer cell lines (Kim *et al*, 2019). Considering the mitochondrial location for PUMA and its recently described role in mitochondrial metabolism in cancer cell lines, we wanted to explore potential mitochondrial functions regulated by PUMA.

**Figure 8.**
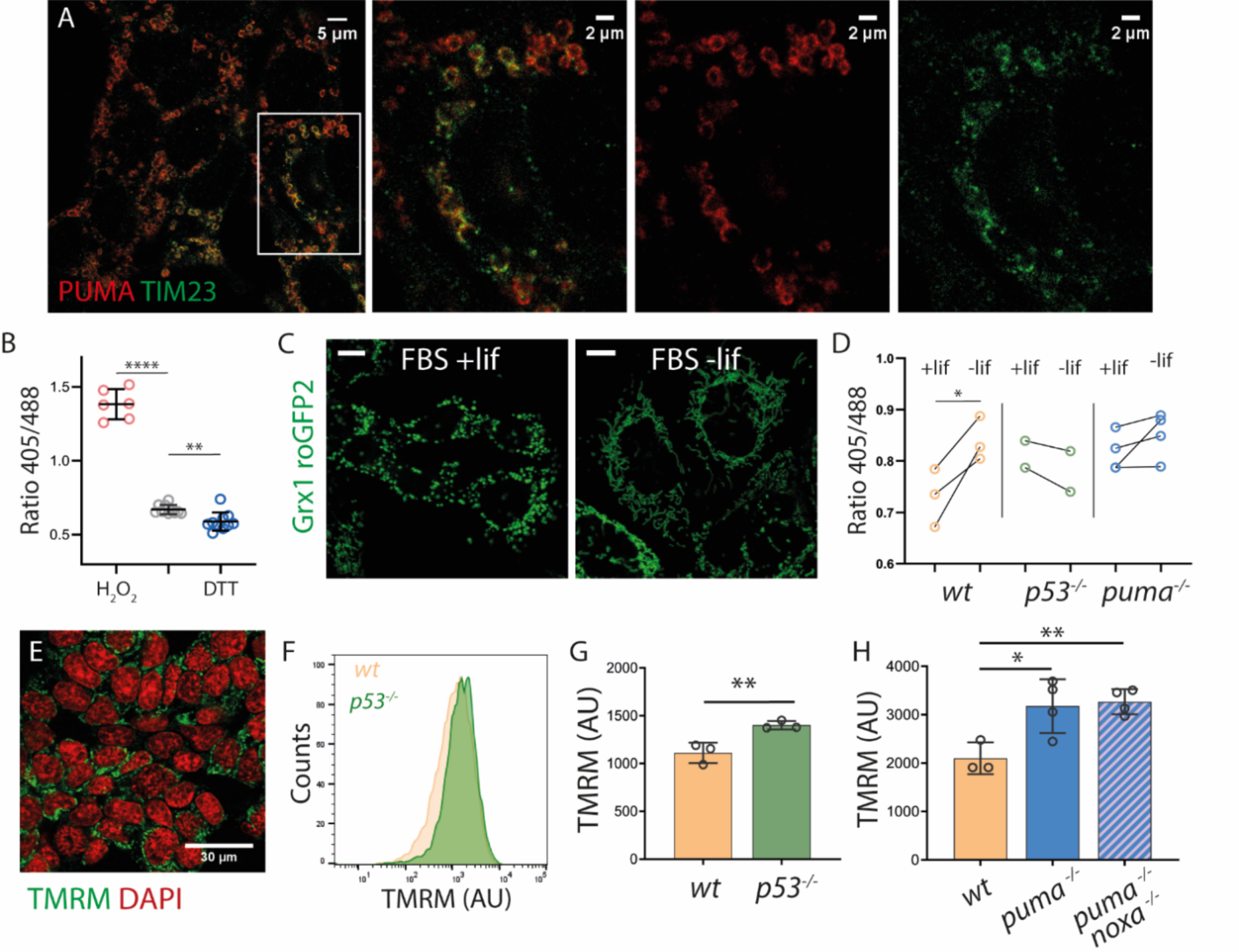
P53 and PUMA regulate mitochondrial membrane potential and oxidative status. **A.** STED-Super-resolution microscopy images showing PUMA and mitochondrial marker TIM23 levels. **B.** 405/488 emission of Grx1-roGFP_2_ expressing ES cells in control conditions (central column) or after the exposure to H_2_O_2_ or DTT. **C.** Confocal images of *wt* cells in conventional and differentiating conditions. **D.** 405/488 emission of *wt*, *p53^-/-^*and *puma^-/-^* Grx1-roGFP_2_ expressing cells in conventional and differentiating conditions. **E.** Confocal capture of TMRM in ES cells. **F.** Histogram of TMRM levels in *wt* and *p53^-/-^* populations. **G, H.** TMRM levels of *wt*, *p53^-/-^, puma^-/-^* or the double knockout *puma^-/-^ noxa^-/-^*cells.

To do so, we first analyzed the mitochondrial REDOX status, as it can vary according to mitochondrial activity (Handy & Loscalzo, 2012). We took advantage of the mitochondrial ratiometric reporter protein Grx1-roGFP_2_, which consists of a modified version of the GFP (roGFP) that adopts an oxidized and a reduced conformation (Østergaard, 2001). When reduced, it has a maximum excitation at λ=488nm and when oxidized, it switches to λ=405nm (Fig. S8A). Thus, the emission intensity when the roGFP is excited at 405 or 488nm varies along with the REDOX status, which is monitored by the emission ratio after excitation at 405 and 488nm. For these experiments, we used a version of the roGFP that is translocated to the mitochondria fused to the glutaredoxin-1 (Grx1) to facilitate the real-time REDOX status of the mitochondria (Gutscher *et al*, 2008) (Fig S8A). First, we checked that the reporter was efficiently oxidized and reduced by using H_2_O_2_ and dithiothreitol (DTT) (Fig. 8B). This showed that ES cells exhibit a REDOX status close to maximal reduction produced by exposure to DTT. Additionally, we found that changes in REDOX status were quick, reaching a maximum oxidation in approximately 7 seconds upon H_2_O_2_ exposure (Fig. S8B, C). Using this tool, we found that the REDOX status of ESC mitochondria did not correlate with PUMA or MYC levels (Fig. S8D, E).

We then studied how the REDOX status varies during pluripotency evolution towards differentiation (Fig. 8C). We found that *wt* cell mitochondria changed to a more oxidized status during differentiation. In contrast, neither *puma^-/-^* nor *p53^-/-^* cells showed this change (Fig 8D). Next, we evaluated the mitochondrial membrane potential (Δψm) as an indicator of differential mitochondrial activity by using TMRM. We found that in the absence of either P53, PUMA or both PUMA and NOXA, cells show an increase in mitochondrial membrane potential (Fig 8 E-H). These results indicate that the activity of the P53-Puma-Noxa pathway regulates mitochondrial activity reducing the transmembrane potential and promoting an increase in mitochondrial oxidation status associated with differentiation.

## DISCUSSION

Cell Competition is a mechanism based on the elimination of viable suboptimal cells, which selects for the fittest cells. It is considered a quality control system conserved and extended in metazoans that may play fundamental roles during development, aging and homeostasis as well as during disease (Merino *et al*, 2015; Coelho *et al*, 2018; Clavería & Torres, 2015; Levayer & Moreno, 2013). This mechanism can function as a tumor suppressor mechanism but can also be adopted by tumour cells to promote their own expansion at the expense of neighboring normal cells (Kon & Fujita, 2021; Morata & Calleja, 2020).

In particular, during the early mouse embryo, CC selects the pool of pluripotent cells that eventually will give rise to the new individual. We know different aspects related to the function of Pluripotent Cell Competition. For examples, it selects the cells with anabolic activity driven by a YAP-MYC pathway (Clavería et al, 2013; Hashimoto & Sasaki, 2019) while removes those cells prematurely differentiating (Díaz-Díaz *et al*, 2017), defective in growth signal detection (Sancho *et al*, 2013), aneuploid (Sancho *et al*, 2013; Singla *et al*, 2020) and those exhibiting a mitochondrial stress signature (Lima *et al*, 2021). Indeed, cell stress constitutes a pivotal aspect in CC, being described in multiple tissues and CC models (Lima *et al*, 2021; Kucinski *et al*, 2017; Wagstaff *et al*, 2013; Baumgartner *et al*, 2021).

In an effort to better understand how fitness is regulated and components that define the “loser status” in Pluripotent CC, we identified the P53 pathway. P53 is a master regulator and reporter of cell stress and is extensively described in mammalian Cell competition (Bondar & Medzhitov, 2010; Marusyk *et al*, 2010; Zhang *et al*, 2017; Wagstaff *et al*, 2016; Díaz-Díaz *et al*, 2017; Fernandez-Antoran *et al*, 2019; Bowling *et al*, 2018; Montero *et al*, 2022).

Downstream P53 pathway, we have identified different candidate genes which can exert a role in the execution of loser cell death and defining fitness (Fig 2B). Here, we have found different elements related to activation of autophagy/mitophagy such as *trp53inp1* or the mTOR inhibitor *ddit4*, Ca^+2^ regulation in endoplasmic reticulum (*perp*) or apoptosis (BCL-2 proteins) (Fig. S11A). We have confirmed that P53 has a role inducing CC and we have reported for the first time that at least some members of the BCL-2 family in addition to their role in apoptosis execution, they show constitutive expression in pluripotent cells with expression levels correlating inversely with competitive fitness. Regarding the mTOR pathway targets, their role has been reported in pluripotent CC (Bowling *et al*, 2018), however, the specific role of autophagy or the endoplasmic reticulum dynamics and their relationship with Pluripotent CC remain to be explored.

Regarding the mechanism by which P53 and BCL2 protein PUMA and NOXA regulate pluripotent cell competitive ability, we have shown that PUMA is strongly localized to mitochondria and its activity reduces the mitochondrial membrane potential and increases mitochondrial oxidative environment. A decrease in the mitochondrial membrane potential has been reported in models of CC in loser BMP mutant cells (*bmpr1^-/-^)* or tetraploid cells (4n) and fission mutant (*mfn2^-/-^, drp1^-/-^)*. Knocking out *p53* in *bmpr1^-/-^* increased the membrane potential over the level of the *wt,* indicating that P53 acts downstream BMP deficiency to affect mitochondrial activity (Lima *et al*, 2021). We thus suggest that P53-PUMA-Noxa regulation of cell competitive fitness is mediated by the role of this pathway on mitochondrial activity.

The knowledge on what determines the activity of the P53 pathway in pluripotent cells remains incomplete. Different types of stresses have been associated with CC, such as oxidative or proteotoxic stress in different models (Kucinski *et al*, 2017; Baumgartner *et al*, 2021). We have explored DNA damage and oxidative stress by measuring the presence of double strand breaks (DSBs) and using the molecular probe dihydroethidium (DHE) respectively, but we did not identify a correlation between these types of stress and MYC levels in ES cells (Fig. S9). Apart from stress, we have shown that P53-PUMA and MYC are regulated by the pluripotency status. Although we did not explore in detail whether P53-PUMA can exert a role in pluripotency, P53 has been associated with ESC differentiation (Lin *et al*, 2005; Jain *et al*, 2012; Zhang *et al*, 2014; Abdelalim & Tooyama, 2014; Fu *et al*, 2020; Jain & Barton, 2018). We found that in the absence of *p53*, the ESCs showed higher levels of p-ERK. This could be interpreted as a direct effect of P53 preventing ESC differentiation, thereby promotes the naïve pluripotent state. Since in these culture conditions naïve cells tend to kill primed cells by cell competition (Díaz-Díaz *et al*, 2017) a possible mechanism for the increase in pERK would be the inhibition of primed-cell death, which would lead to their accumulation. Against this idea, in the same test, elimination of *puma* or *noxa* does not change the pERK pattern, suggesting that these death pathways are not involved in the role of P53 on ES cell differentiation.

The constitutive expression of PUMA in early mouse embryos appears exclusively related to pluripotency, and to the presence of endogenous CC in the early mouse embryo. Indeed, only epiblast cells show this widespread PUMA expression profile and its expression is shut down in gastrulating cells. This change in expression might be related to a shift in its function from a factor that regulates fitness to a direct death inducer factor. In this respect, the recent mitochondrial metabolic role described for PUMA in cancer cells assimilates these to pluripotent cells of the early embryo (Kim *et al*, 2019). A pattern of high expression of PUMA in a heterogeneous pattern therefore appears to be related to pluripotency *in vitro* and *in vivo*.

Interestingly, we have reported that P53 exerts an inhibitory effect over MYC. However, ES cells still exhibit a MYC heterogeneous expression pattern in the absence of P53, indicating that P53 is not solely responsible for the variability in MYC expression in ES cells but there should be other factors regulating MYC expression. Notably, *puma*^-/-^ or the double knockout *puma^-/-^*, *noxa^-/-^*cells do not present changes in MYC expression, indicating that MYC and PUMA/NOXA regulate independent branches of P53-induced CC (Figure S10)

## MATERIALS AND METHODS

### Cell line generation

The *Myc^GFP/GFP^* allele was described in (Huang *et al*, 2008) and the mES cell line carrying the allele was described in (Díaz-Díaz *et al*, 2017). The MYC overexpressing cell line *iMOS^T1-MYC^* was described in (Clavería *et al*, 2013). *Myc^-/-^* ES cell line was generated in our lab by recombining the *Myc^flox^* allele. *P53, puma, Noxa* and *Myc* knockout ES cell lines were generated with CRISPR-CAS9 technology using the *Myc^GFP/GFP^* ES cell line. Two crRNA sequences were employed per gene to generate a deletion in the gene sequence. The web tool CRISPOR (http://crispor.tefor.net/crispor.py) (Concordet & Haeussler, 2018) was used for the design of the crRNAs which are indicated in Table 1. In the case of *p53*, the targeted region included the DNA binding domain, the nuclear localization sequence and the oligomerization domain. Regarding *puma*, the deleted region covers the majority of exons 1 and 2, including the BH3 domain and the Ser96 and Ser106 residues, recently described as key for PUMA-metabolic functions (Kim *et al*, 2019). In the case of *noxa*, the removed region consisted in exons 2 and 3, including BH3-1 and 2 domains. In the case of *Myc*, deletion of the second exon of the gene, including the whole gFP coding region and was deleted.

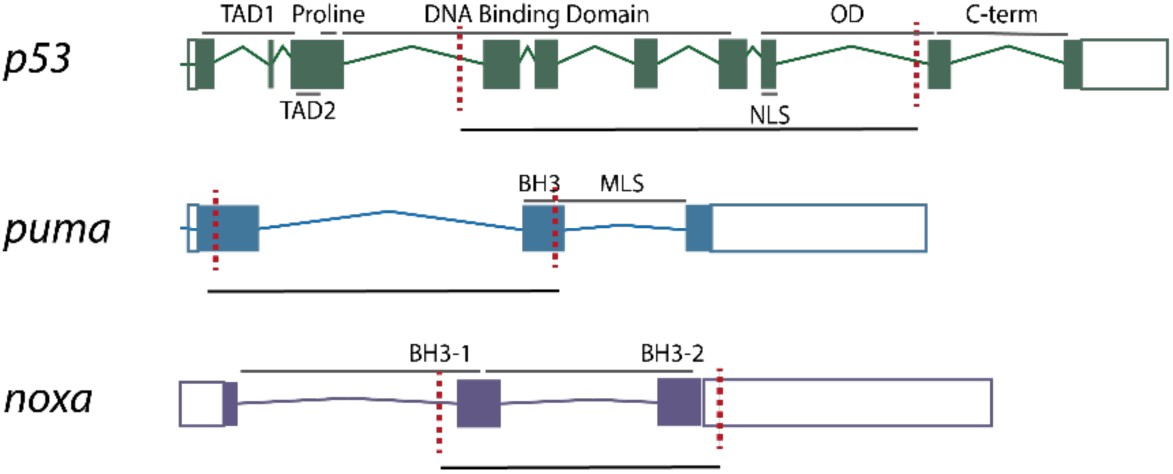

**TABLE 1.**
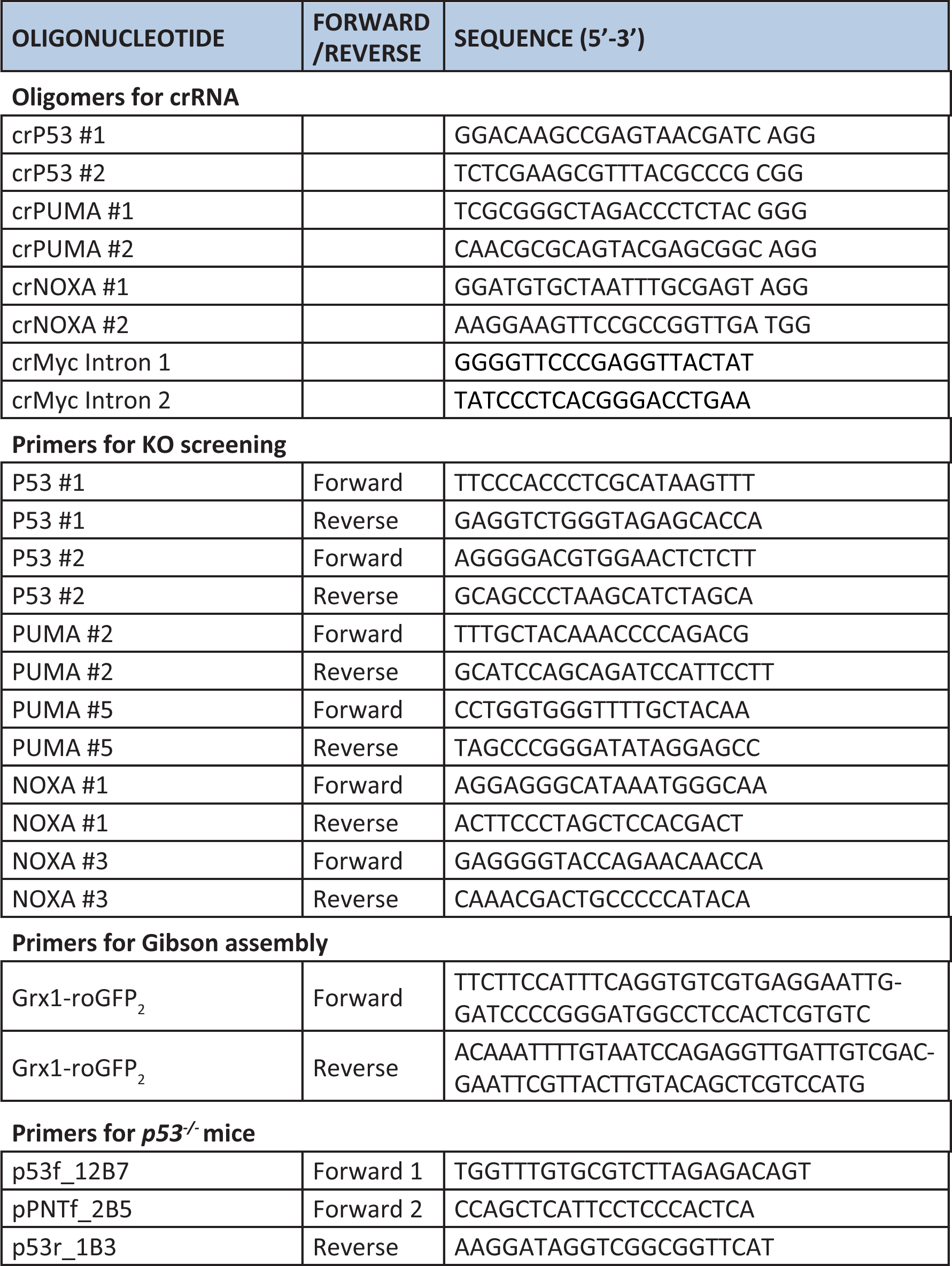
crRNA and primer sequences.

crRNAs and tracrRNA were purchased from IDT while CAS9 protein was expressed and purified by the Pluripotency Cell Unit at CNIC. To generate each knockout cell line, 2×10^6^ Myc^GFP/GFP^ cells were electroporated together with the ribonucleoprotein (RNP) complex, formed by the guide RNA (crRNA + tracrRNA) and the CAS9. Cells were electroporated with the Neon Transfection System (Thermo Fisher Scientific). A *tdtomato*-expressing plasmid was used as a reporter, so that 24h after the electroporation, tdTomato positive cells were sorted by FACS. tdTomato expressing cells displayed due to the electroporation, allowing the *tdtomato* plasmid to get into and being expressed, being more likely that the RNP complex entered into these cells. Then, individual cells were expanded into single colonies and knockouts clones were screened by PCR.

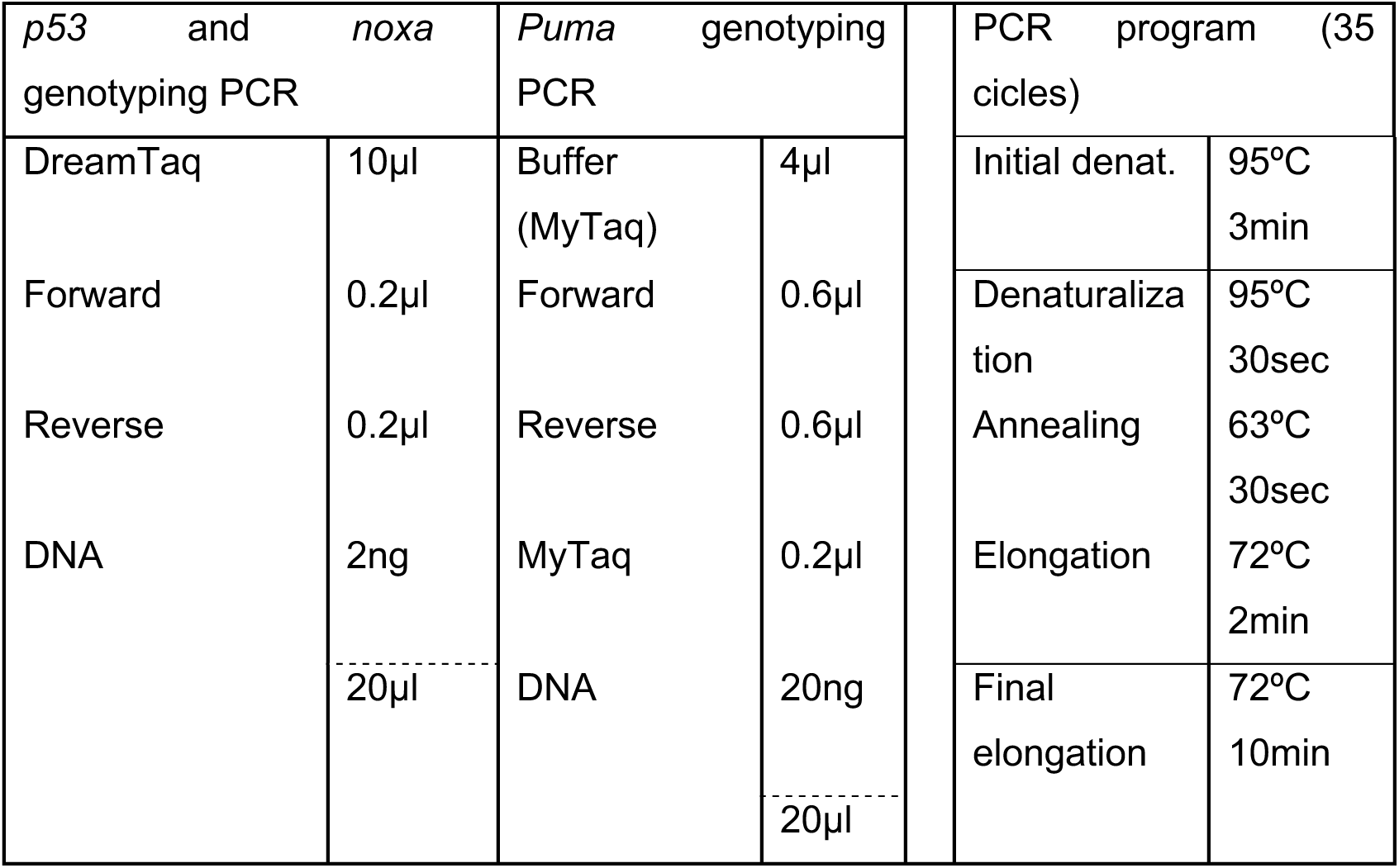

ES cell lines expressing the *Grx1-roGFP2* REDOX reporter were generated by lentiviral transduction. First, we cloned the *Grx1-roGFP2* sequence from the “pLPCX mito Grx1-roGFP2” plasmid into a lentiviral vector under the EF1 promoter “pCDH-EF1” using Gibson Assembly (NEB) (primers described in Table 1). pLPCX mito Grx1-roGFP2 was a gift from Tobias Dick (Addgene plasmid #64977), pCDH-EF1 was a gift from Kazuhiro Oka (Addgene plasmid # 72266). The final sequence was then packaged into lentiviral particles by the Viral Vectors Unit at CNIC. Finally, lentiviral particles were used to transduce 100.000 ESCs with a MOI between 5 and 10 ON in a 24-well plate. After ESCs expansion, GFP positive cells were sorted by FACS.

### Animals

Animals were handled in accordance with CNIC Ethics Committee, Spanish laws and the EU Directive 2010/63/EU for the use of animals in research. All mouse experiments were approved by the CNIC and Universidad Autónoma de Madrid Committees for “Ética y Bienestar Animal” and the area of “Protección Animal” of the Community of Madrid with references PROEX 220/15 and PROEX 144.1/21. Wild-type mice were on a CD1 out-bred genetic background. *p53^-/-^*mice were generated by crossing heterozygous *p53^tm1b/+^* mice, previously described in (http://www.informatics.jax.org/allele/MGI:6120822) (Jacks *et al*, 1994). *p53^tm1b/+^* animals were kindly gifted by Ignacio Flores lab. *P53^tm1b^*animals were genotyped using DreamTaq Green (ThermoFisher) as indicated below and the primers are included in (Table 1).

| *p53* genotyping PCR |  | PCR program (35 cycles) |  |
| --- | --- | --- | --- |
| DreamTaq | 10μl | Initial denat. | 94°C<br>3min |
| Forward 1 | 0.2μl | Denaturalization | 94°C<br>30sec |
| Forward 2 | 0.2μl | Annealing | 60°C<br>30sec |
| Reverse | 2ng | Elongation | 72°C<br>1min |
| DNA | 1μl | Final elongation | 72°C<br>10min |
|  | -----<br>20μl |  |  |

### Embryo retrieval

Midday of the day when vaginal plug was detected was considered gestational day 0.5 (E0.5). Females were culled by CO_2_ inhalation and the uterus was and transferred to DMEM media #41965-039 (ThermoFisher) at 37°C. Embryo extraction at E3.5 was performed by the blastocysts out of the uterus under a dissection scope using a 1ml syringe with a 23-G needle. Blastocysts were transferred to KSOM media MR-101 (Merck) with a mouth pipette. Tyrodés solution #T1788 (Merck) was used during a few seconds to dissolve zona pellucida. Blastocysts were then washed in PBS 1% FBS and fixed in paraformaldehyde (PFA 2%) dissolved in PBS 1% FBS overnight at 4°C. Eventually, blastocysts were washed in PBS + Triton-X100 0,1% 1% FBS. For E5.5-E.7 embryos extraction, working under the scope and using precision forceps (Dumont #55 0.05×0.02mm) (FST), muscular uterine walls were carefully ripped. After that, both the decidual layer and the Reichert’s membrane were removed and embryos were fixed in paraformaldehyde (PFA) (Merck) 2% in PBS overnight at 4°C. After fixation, embryos were washed in PBS several times.

### Cell culture

Mouse embryonic fibroblasts (MEFs) were used as feeder cells for the ESCs. Fibroblasts were initially extracted from E10.5 CD1 embryos by the Pluripotent Cell Technology unit at CNIC. 5 million fibroblasts were then plated on a 100mm plate in MEFs medium (described below) and passaged to three 150mm-plate 48h later. After 3-4 days, MEFs were inactivated using mitomycin C #M4287 (Sigma) during 2.5h and washed 3 times using PBS. Finally, MEFs were trypsinized (Trypsin-EDTA 10x, Gibco) and frozen (1,2 million cells in 1ml of freezing medium). Upon inactivation, MEFs were plated on 0,1%-gelatin coated plates. For cell expansion, 1 million mouse embryonic stem cells (mESCs) were plated on a 35mm-plated previously covered by inactivated MEFs. After 48 hours, cells were passaged on a 100mm-plate covered by inactivated MEFs and cells were trypsinized and frozen two days later (1.2 million cells in 1ml volume per cryovial). To perform experiments, ESCs were thawed on a 35mm-plate covered by inactivated MEFs. After 2 days, MEFs depletion is performed and ESCs are transferred to a 0,1%-gelatin coated 35mm-plate. MEFs depletion is based on the different attachment of MEFs and ESCs, so after cells are trypsinized, MEFs attach in 1-2hours, while ESCs can be transferred to a new plate. 0.7 million cells were cultured on a 35mm-plate and 0.18 million on a 12-well plate. For freezing, cells were resuspended in MEFs medium and freezing medium was carefully added in a final proportion 1:1 and aliquot into cryovials. Cryovials are kept in a freezing container (Nalgene®) at −80°C and transferred into liquid nitrogen after 24h.

### Medium composition

MEF medium contains high glucose DMEM (#41965, LifeTech), 15% Fetal Bovine Serum (FBS), 1% sodium pyruvate (#11360070, ThermoFisher), 1% Penicillin/Streptomycin (10,000U/ml; 100x), 0,1% 2-beta-mercaptoethanol (50mM). mESC medium (FBS+LIF) consisted of MEF medium plus 1% non-essential amino acids (100x) and 2x LIF (leukemia inhibitory factor). LIF was provided by the PCT unit and used 250x. LIF was removed to induce mESC differentiation. FBS was substituted by KnockOut™ serum replacement (here referred as SR) (#10828, Invitrogen) to get SR+LIF medium. Finally, the inhibitors CHIR99021 (#04-0004-02, Stemgent) and PD0325901 (#04-0006, Stemgent) were added at 0.1μM and 0.3μM to the FBS+LIF medium to obtain “2i medium”.

### Competition assays

0.18 million cells were plated in co-culture or separated using 12-well plates and FBS medium without LIF. At every time point, cells were trypsinized and counted using a Neubauer chamber (Sigma-Aldrich). The percentage of fluorescent and non-fluorescent cells in co-culture was determined by flow cytometry.

### RT-PCR

For RNA extraction, 4 million cells per ml were resuspended in TRI Reagent (Invitrogen) for 5 min at room temperature (RT). Afterward, ethanol was added in a 1:1 volume proportion and vortex. “Direct-zol RNA Miniprep kit” (R2051, Zymo Research) was used to extract the RNA according to the manufacturers. Finally, RNA was stored at −80°C. To perform the cDNA reverse transcription, we used 1μg of RNA and the “High Capacity cDNA Reverse Transcription’’ Kit (4368814, ThermoFisher). Eventually, we performed a qPCR using “Sybr Green’’ (#4472903, Invitrogen). The primers for the qPCR reaction were purchased from “KiCqStart® SYBR® Green Primers” (Sigma-Aldrich) and the gene *gadph* was chosen as a control.

### Whole-mount Embryo Immunofluorescence

E3.5 whole-mount immunostaining was performed using 4-well plates and a mouth pipette. Triton X-100 0,1% and FBS 1% was added to the PBS and the blocking solutions to avoid blastocyst getting attached to the plate. E5.0-E7.5 embryo immunostaining was performed using 35mm plates and/or round bottom 2 ml Eppendorf tubes using a micropipette with end-cut tips to avoid excessive pressure when transferring the embryos. Both E3.5 and E5.0-E7.5 embryos were permeabilized using 0,5% PBT (PBS + Triton X-100 0,5%) for 20 min at RT. Embryos were washed in PBT 0,1% and blocked using 10% goat serum (Gibco-BRL Life-Technologies) in 0,3% PBT during 1 hour at RT. Embryos were incubated with primary antibodies overnight using blocking solution at 4°C. Embryos were then washed several times with PBT 0,1% and incubated with the secondary antibodies, Wheat Germ Agglutinin (1:500) (ThermoFisher) for plasma membrane staining and/or DAPI (1:1000) using blocking solution for 1 hour at RT. Finally, embryos were washed several times and embedded in mounting media. To avoid the embryos to collapse due to the different density between 0,1% PBT solution and mounting media, the mounting media was diluted in serial dilutions using 0,1% PBT, (25, 50, 80 and 100% mounting media) and the embryos were transferred through the different dilutions. For E.5.0-E7.5 embryos, we used VectaShield mounting media (H-1000-10, Vector Laboratories) while, for E3.5, liquid Abberior (MM-2013, Abberior) mounting media was used.

### mESCs Immunofluorescence

0.18-0.25 million ES cells were plated on 35mm-glass bottom dishes (MatTek), previously coated using human fibronectin (#354008, Corning) according to the manufacturers ON at RT. Two days after plating, cells were washed with PBS and fixed with PFA 2% ON at 4°C. For mESCs, a similar procedure than with embryos was carried out, but permeabilization was reduced to 10 minutes and Vectashield was used as mounting media. Primary and secondary antibodies were incubated in a 100μl volume. To prevent evaporation during the primary antibody overnight incubation, plates were stored inside a humidity chamber. For 594-conjugated cleaved-CASP3 (#8172, Cell Signalling) immunostaing, we proceed as described by the manufacturers (Protocol Id: 182) (https://www.cellsignal.com/products/antibody-conjugates/cleaved-caspase-3-asp175-d3e9-rabbit-mab-alexa-fluor-594-conjugate/8172). To induce activation of P53, ESCs were exposed to etoposide (Sigma) 40μM during 10 hours or Nutlin-3 (BioVision) 5-30μM during 12 hours previous to fixed. For immunostaining of ESCs in suspension, ES cells were trypsinized and fixed with PFA 2% in PBS during 1 hour at 4°C. Saponin substituted Triton-X as a permeabilizing agent and is added at 0,1% in all solutions. After the immunostaining, ESCs were washed and resuspended in 200μl of PBS and analyzed by flow cytometry. Primary antibodies are summarized in Table 2.

**Table 2.**
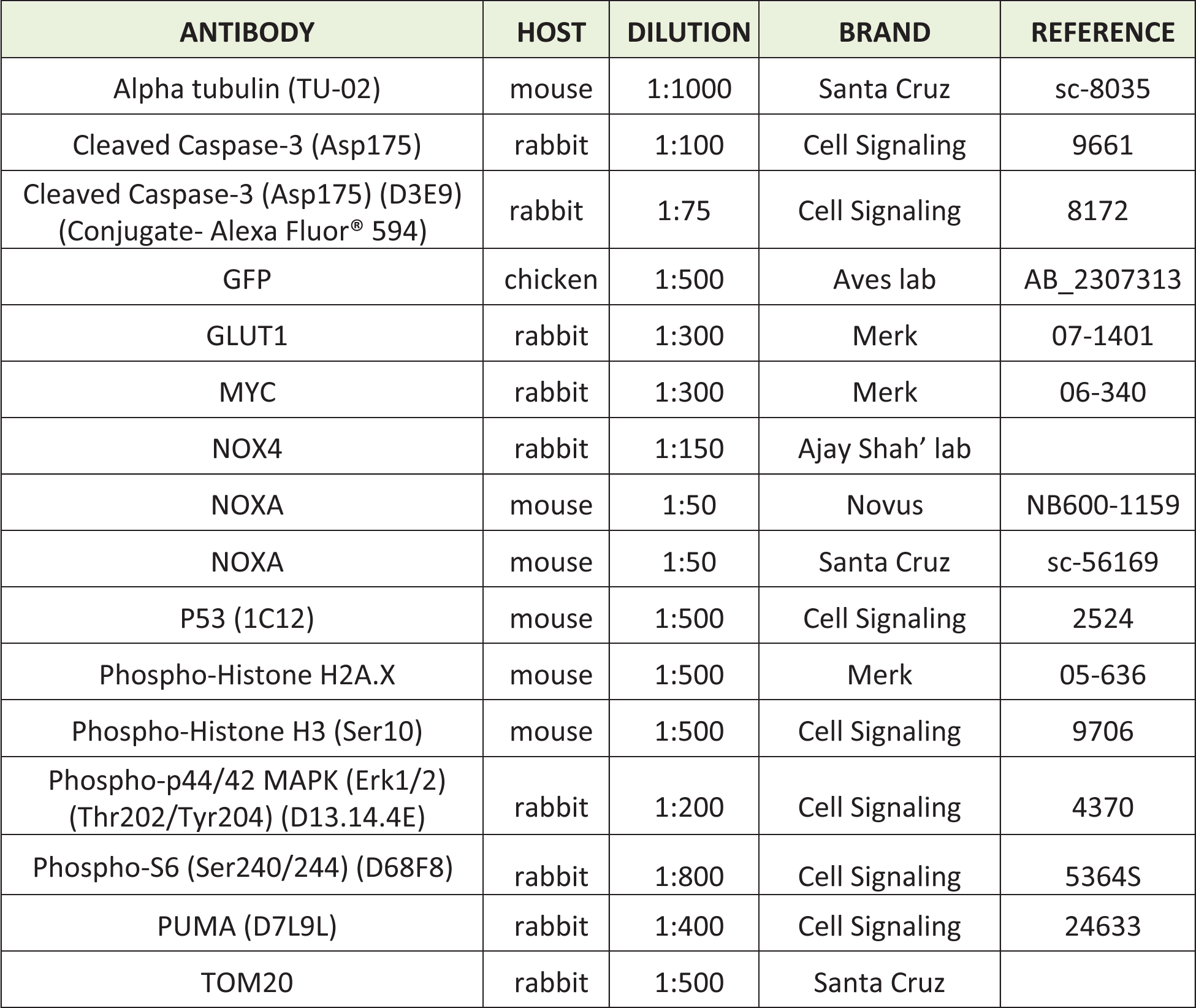
Antibodies.

### ROS Measurement

DHE (Invitrogen) was used for H_2_O_2_ and O_2_^.-^ measurement. ESCs were incubated with DHE 2μM for 15 min at 37°C. Cells were washed and analyzed “in live” by confocal microscopy. H_2_O_2_ 0,2% was used as a positive control while “cold-ice incubation” with the DHE was used as a negative control.

### Mitochondrial Membrane Potential

TMRM (D-1168, Invitrogen) was used to measure membrane potential. Cells were incubated with TMRM 25nM for 20 min at 37°C. Cells were washed with PBS and analyzed “in live” by confocal microscopy. Cells were exposed to oligomycin (Sigma-Aldrich) 2µM for 3 hours previous to the TMRM treatment as a positive control while FCCP (Sigma-Aldrich) 50µM for 30 min was used as a negative control.

### Apoptosis Measurement

FLICA™ 660 Caspase-3/7 (BIORAD) was used to measure apoptosis according to manufacturers.

### Cell Cycle

For Cell Cycle analysis, 0.2 million cells were trypsinized and resuspended drop-by-drop with 500μl of ethanol 70% at −20°C and stored for 24h at −20°C. Subsequently, cells were washed in PBS twice and resuspended in 200μl of PBS containing propidium iodide at 50μg/ml. Cells were analyzed by flow cytometry. Dean-Jett-Fox model was used for this analysis.

### Mitochondrial REDOX status measurement

ESCs stably expressing Grx1-roGFP2 were used to study the mitochondrial REDOX status. When cells were fixed, N-ethylmaleimide (NEM) (Sigma) 20mM was used 5min previous to fixation to protect the cells against the thiol oxidation mediated by paraformaldehyde fixation (Albrecht *et al*, 2011). Exposure to H2O2 100μM or DTT 20mM was used to promote maximum and minimum oxidation.

### Immunoblot

Cells were lysed with RIPA buffer containing (25x) protease inhibitor (Roche) for 30 min at 4°C. Approximately 1.5 million cells were lysed at a concentration of 3 million cells per ml. Protein concentration was measured using BSA (Sigma-Aldrich) serial dilutions and the “DCTM Protein Assay” kit (BioRad). Absorbance was measured at 690 nm using a microplate spectrophotometer. Proteins were separated via 12% SDS-PAGE under reducing conditions and transferred to a polyvinylidene difluoride (PVDF) membrane using “Wed blotting system” (BioRad). After incubation with primary and secondary antibodies, protein signal was detected via chemiluminescence using the “Pierce ECL Western Blotting Substrate” kit (Thermo-Fisher).

### Equipment

Confocal microscopy was performed using a Leica TCS SP8 coupled to a DMi8 inverted confocal microscope Navigator module equipped with white light laser. A HC PL Apo CS2 40x/1.3 oil objective and 1024×1024 pixels, A.U. set to 1 were commonly used. For Super-resolution microscopy, a Leica gated STED-3X-WLL SP8 and a HC PL APO CS2 100x/1.40 oil objective was used. Alexa Fluor 514 and Alexa Fluor 568 secondary antibodies were used for this technique. Flow Cytometry was performed using a BD LSRFortessa Special Order Research Product (laser wavelengths 405, 488, 561, 633). Additionally, ESCs were sorted using a BD FACSAriaTM II and Synergy 4L cell sorter.

### Image Analysis

Confocal images were analyzed using FIJI (Schindelin *et al*, 2012) (https://imagej.net/Fiji). For nuclear segmentation, we took advantage of DAPI/TO-PRO-3TM staining. Nuclei masks were created applying the “default Threshold tool” and manually corrected to ensure that segmented objects correspond to individual cells and discard mis-located, apoptotic or mitotic cells. Finally “Analyzed particle” tool was used to identify the regions of interest (ROIs).

For cytoplasmic segmentation, we used the WGA membrane staining. We applied a “Gaussian Blur” filter (scaled units 2) and the “Find Maxima” tool (configuration: output type, segmented particles; light background) to create a mask. Afterwards, manual correction was performed to ensure that segmented objects correspond to individual cells, and we applied “Analyzed particles” to identify the ROIs. Finally, we subtracted the nuclear area (obtained as described above) to the whole cell region. To couple the cellular and nuclear ROIs from the same cell, a macro was designed together with the CNIC Microscopy Unit (Anexo 1).

To quantify nuclear foci, ROIs corresponding to a nucleus were selected and processed using the “Find maxima” tool (output=count). This process was automated by establishing a running a macro with a loop (Anexo 1).

To study PUMA and MYC correlation in E6.5 mouse embryos, MYC levels were normalized per Z slice and embryo to avoid depth-dependent loss of signal and variation among different embryos. Statistical analysis was performed using linear mixed models using lme4 R library, p value=6.81×10^-10^. This type of analysis was also performed for the P53-PUMA correlation at E3.5, p=0.105 and P53-MYC correlation at E3.5. Embryo was set as a random variable and either P53, PUMA or MYC-classification and Z-position as covariates to simultaneously adjust for the two factors. Coefficients represent either the quantitative increase in the response variable (log2(MYC)) per unit increase in the independent variables (either log2(P53) or PUMA-classification variables) and their associated p-values show the significance of such coefficients under the null hypothesis of them being 0.

### Statistical analysis

Parametric T student test was performed to compare two groups of data. For comparisons with more than two groups of data One –way ANOVA multiple comparison was used. One-sample test (Wilcoxon test) was used to compare a group of data with a hypothetical mean. Comparison and graphs were made with Graph Pad Prism 8.4.3 statistical analysis software. Adjusted values of P<0,05 were considered statistically significant.

### RNAseq Meta-Analysis

RNAseq data for meta-analysis were initially described in (Díaz-Díaz *et al*, 2017). For the enrichment analysis, we used the web server gProfiler (version e108_eg55_p17_9f356ae) with g:SCS multiple testing correction method applying significance threshold of 0.05 (Raudvere *et al*, 2019). For the analysis, we previously selected those genes upregulated in MYC-LOW cells and we filtered those genes with more than 3 reads after normalization. For the selection of candidate genes from our RNAseq data related to the P53 pathway and associated to apoptosis/cell stress, we took advantage of Gene Ontology and GO Annotation, using the QuickGO (https://www.ebi.ac.uk/QuickGO/) developed by the EMBL’s European Bioinformatics Institute (EMBL-EBI).

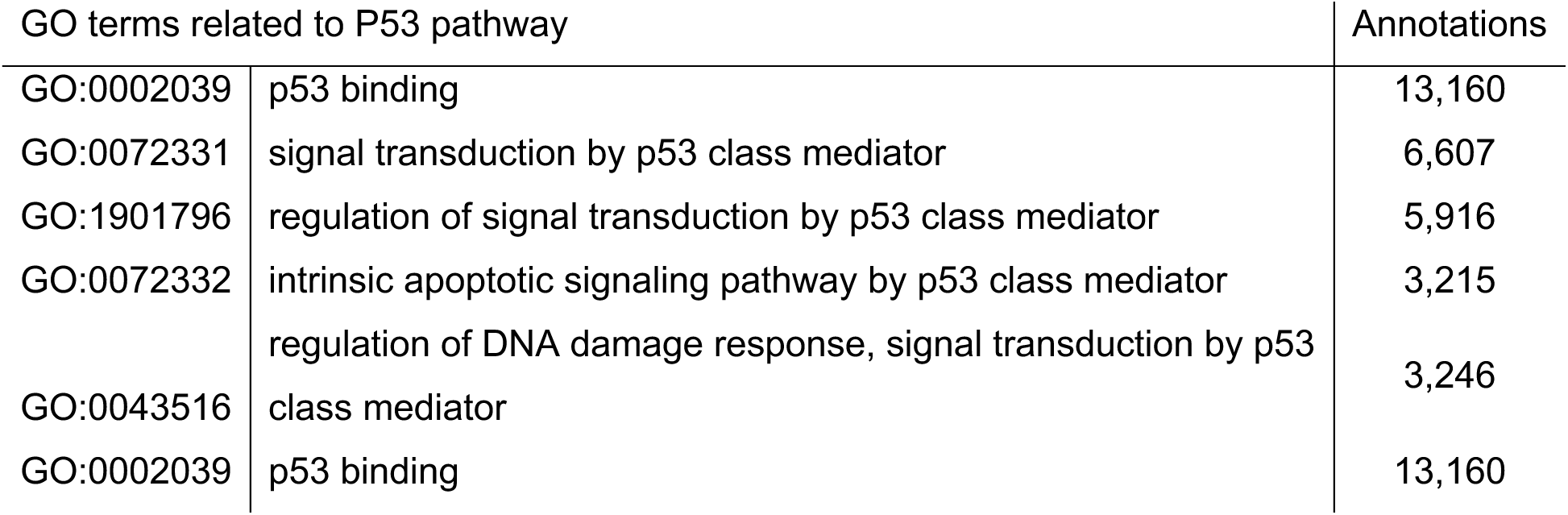

| GO terms related to apoptosis |  | Annotations |
| --- | --- | --- |
| GO:0042981 | regulation of apoptotic process | 213,351 |
| GO:1900119 | positive regulation of execution phase of apoptosis | 463 |
| GO:1900118 | negative regulation of execution phase of apoptosis | 563 |
| GO:0097190 | apoptotic signaling pathway | 28,464 |
| GO:0097194 | execution phase of apoptosis | 4,864 |

For the Volcano plot representation, we used the web app VolcaNoseR tool (Goedhart & Luijsterburg, 2020) (https://huygens.science.uva.nl/VolcaNoseR2/) maintained by Joachim Goedhart and Martijn Luijsterburg. We filtered the genes with more than 3 reads and discarded those without a term in the MGI (Mouse Genome Informatics) database and those with an annotation starting by Gm or ending in Rik. The genes are represented by the log_2_ Fold change and the -log_10_ AdjPValue.

## ACKNOWLEDGEMENTS

We thank members of the Torres laboratory for fruitful discussions and advice on this work. We thank Andrea Curtabbi, Raquel Justo and Jose Antonio Enríquez for generous advice on this work and Ignacio Flores for the P53 mutant mice. We thank the CNIC cytometry, viral vectors and compared medicine technical units for their support to this work. We thank the CNIC Advanced Microscopy Unit for advice on confocal super-resolution microscopy,

---

**Figure S1.**
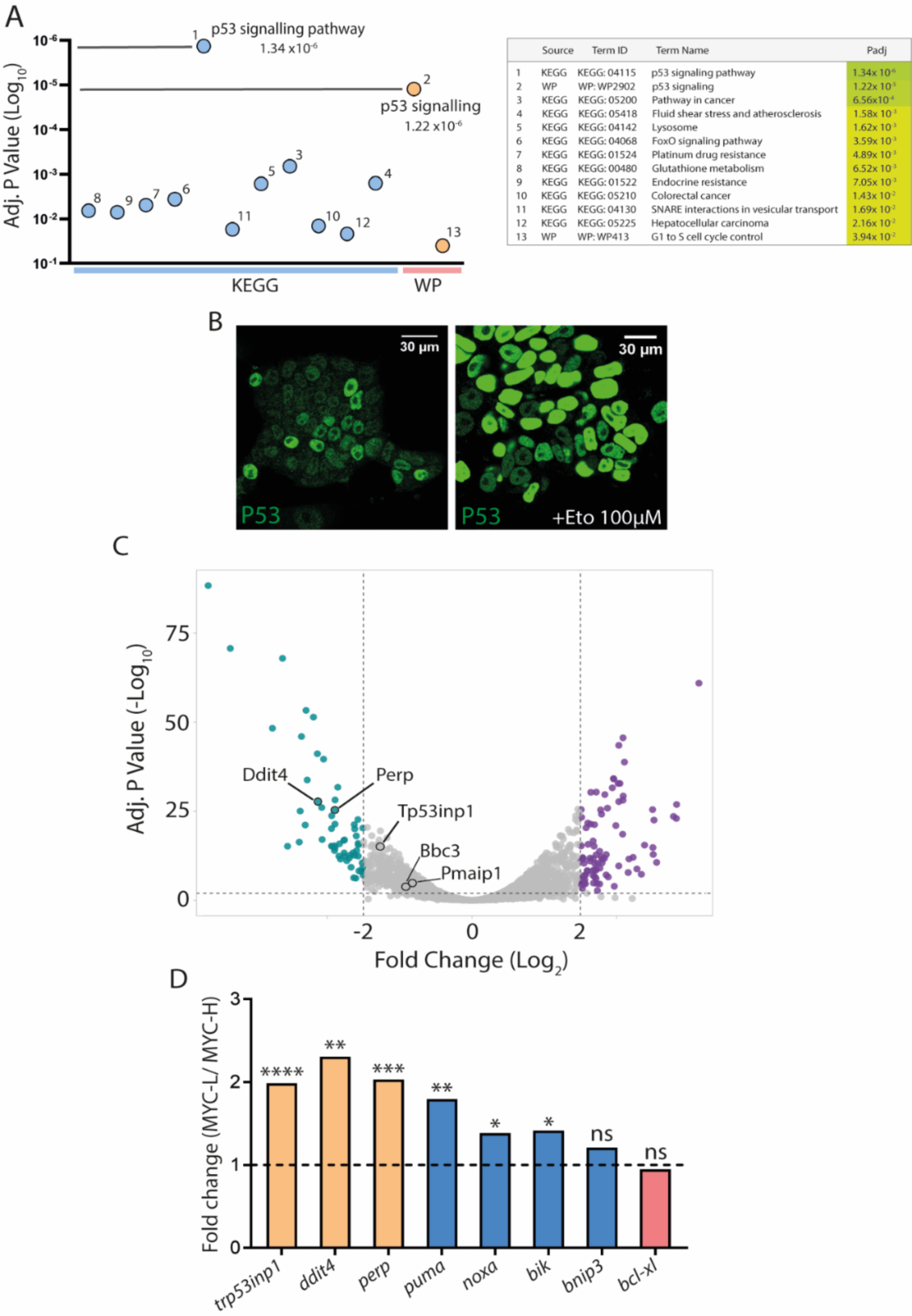
Pathway enrichment analysis and identification of candidate genes downstream P53. **A.** Dot plot and table showing the most enriched pathways and terms associated with our RNAseq data. **B.** Confocal images showing P53 expression in normal conditions and after the exposure to etoposide. **C.** Volcano plots showing the genes from our RNAseq data. Different candidate genes related to the P53 pathway and apoptosis were highlighted. **D.** Bar graph showing the MYC-low versus MYC-high ratio from a qPCR of the indicated candidate genes.

**Figure S2.**
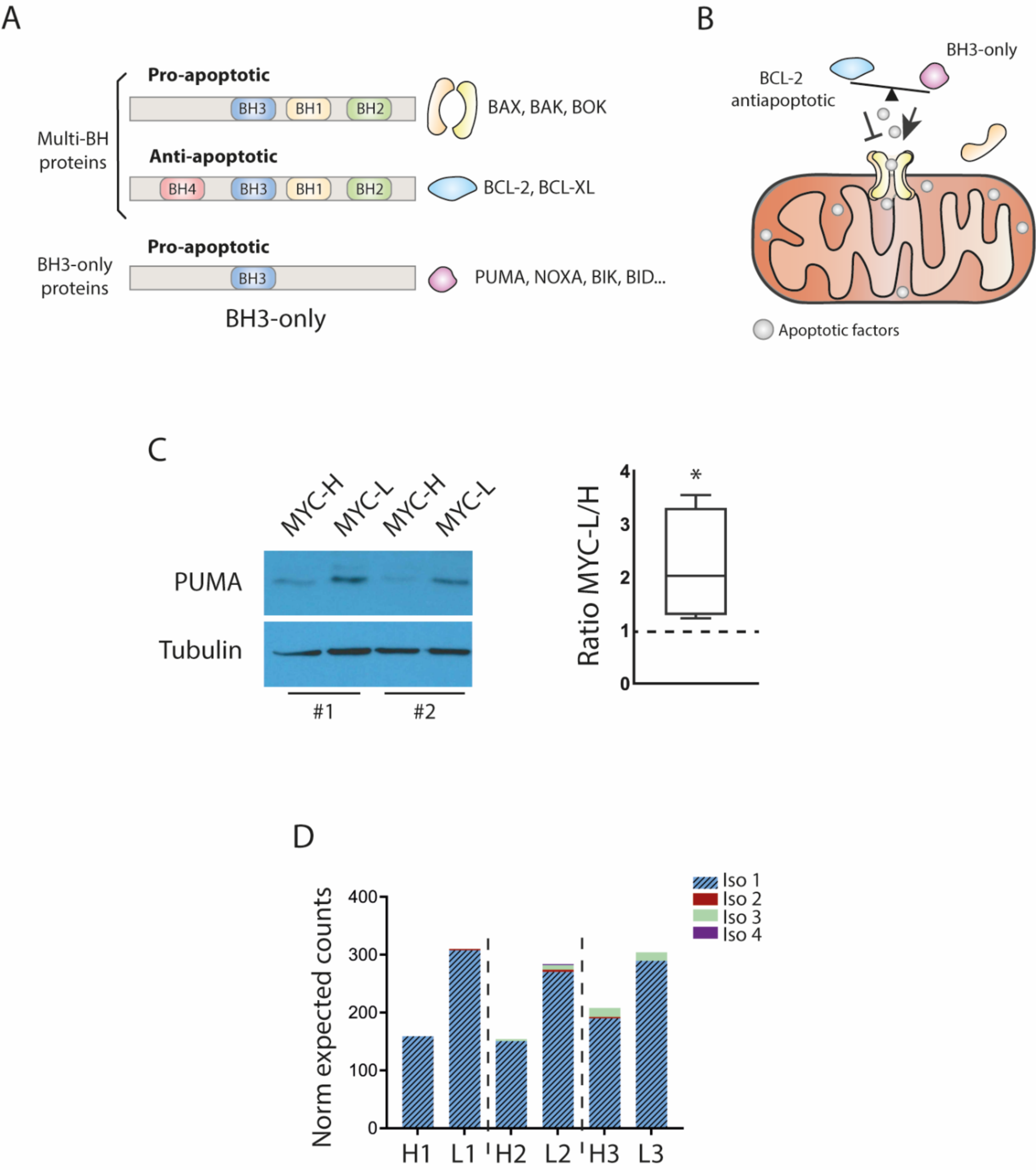
BCL-2 family pro-apoptotic protein PUMA displays a higher expression in MYC-low cells. **A.** Schematic representation of the different BCL-2 protein subfamilies and mechanism of action (**B**). **C.** (left) Western blot of PUMA expression in MYC-H and MYC-L population and quantification (right). **D.** RNAseq data analyses indicating the normalized expected counts of the 4 isoforms of *puma*.

**Figure S3.**
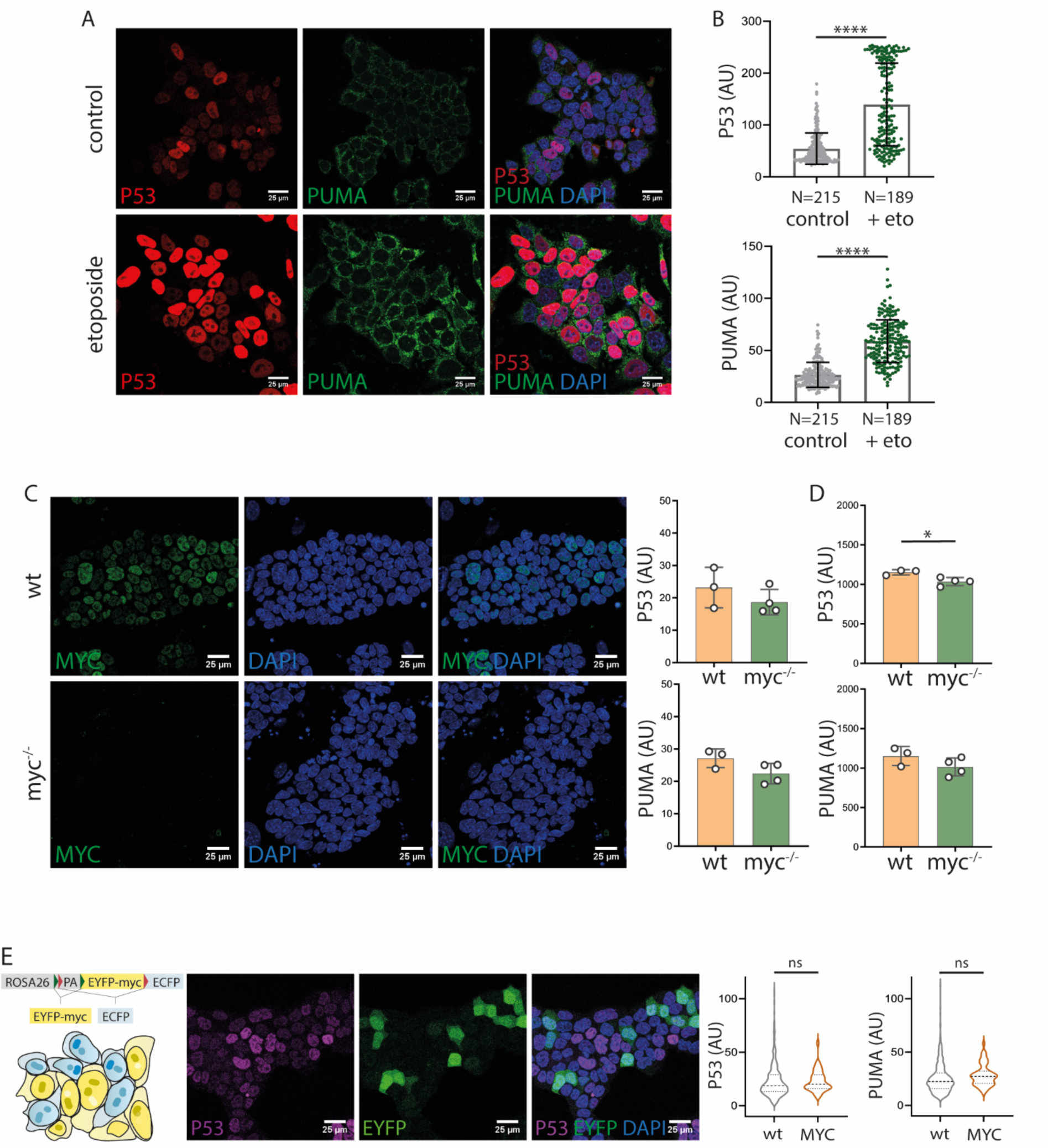
MYC does not regulate P53 or PUMA. **A.** Confocal images showing P53 and PUMA expression with or without etoposide and quantification (**B**). **C.** Confocal captures showing MYC levels in *wt* and *myc^-/-^* cells (left) and quantification of P53 and PUMA levels in *wt* and *myc^-/-^* cells (right). Each dot represent one *wt* or *myc^-/-^* clone. **D.** Bar graph showing the similar experiment than in C but analyzed by flow cytometry. **E.** Schematic representation of the iMOS^MYC^ system (Clavería *et al*, 2013) (left). Confocal images showing P53 and EYFP expression and quantification of P53 and PUMA levels in *wt* cells and cells overexpressing MYC (right).

**Figure S4.**
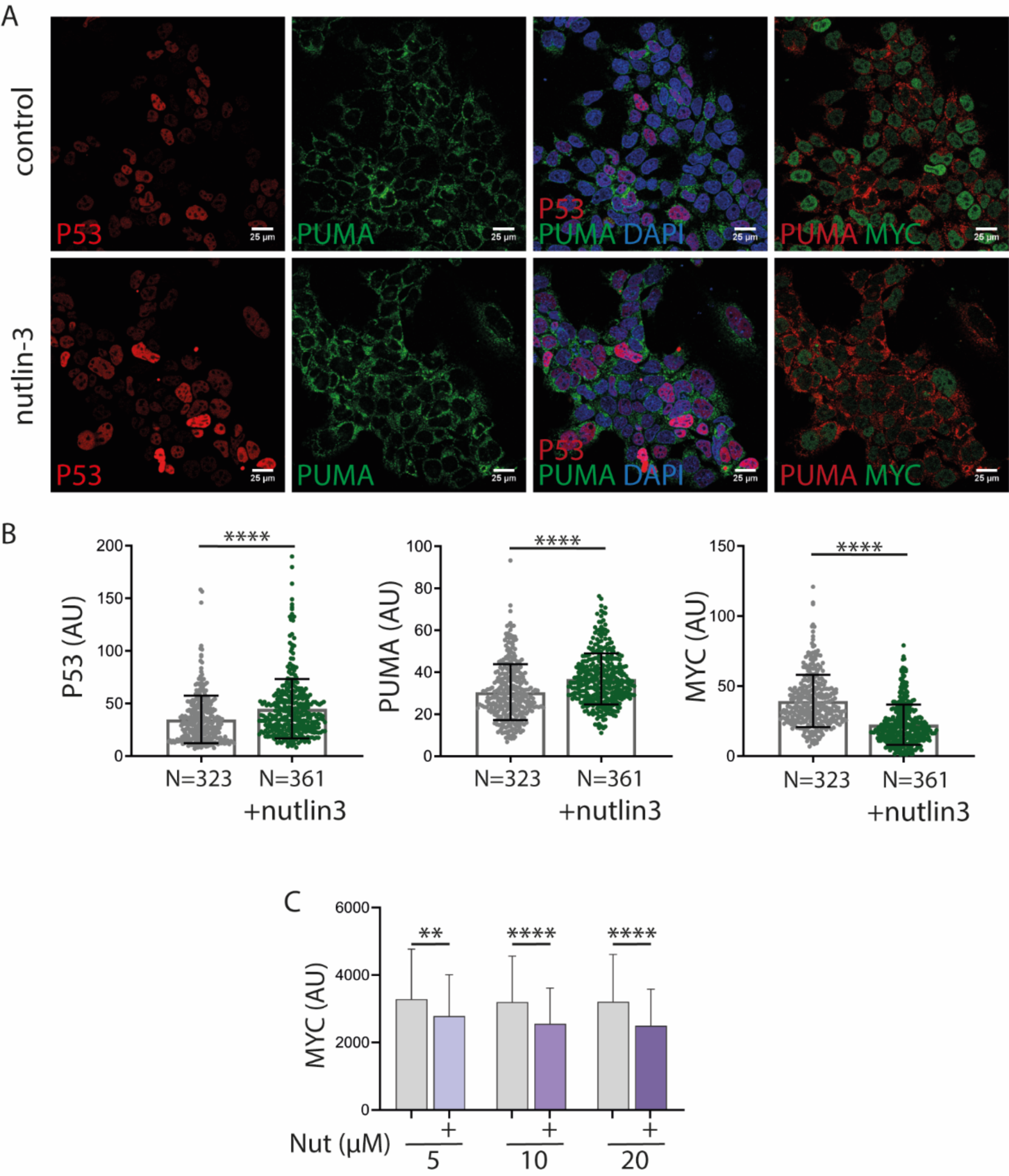
P53 activation using Nutlin3 induces PUMA upregulation and MYC inhibition. **A.** Confocal images showing P53, PUMA and MYC levels in normal conditions and after Nutlin3 treatment and quantification (**B**). **C.** MYC levels upon treatment with different doses of Nutlin3, analyzed by flow cytometry.

**Figure S5.**
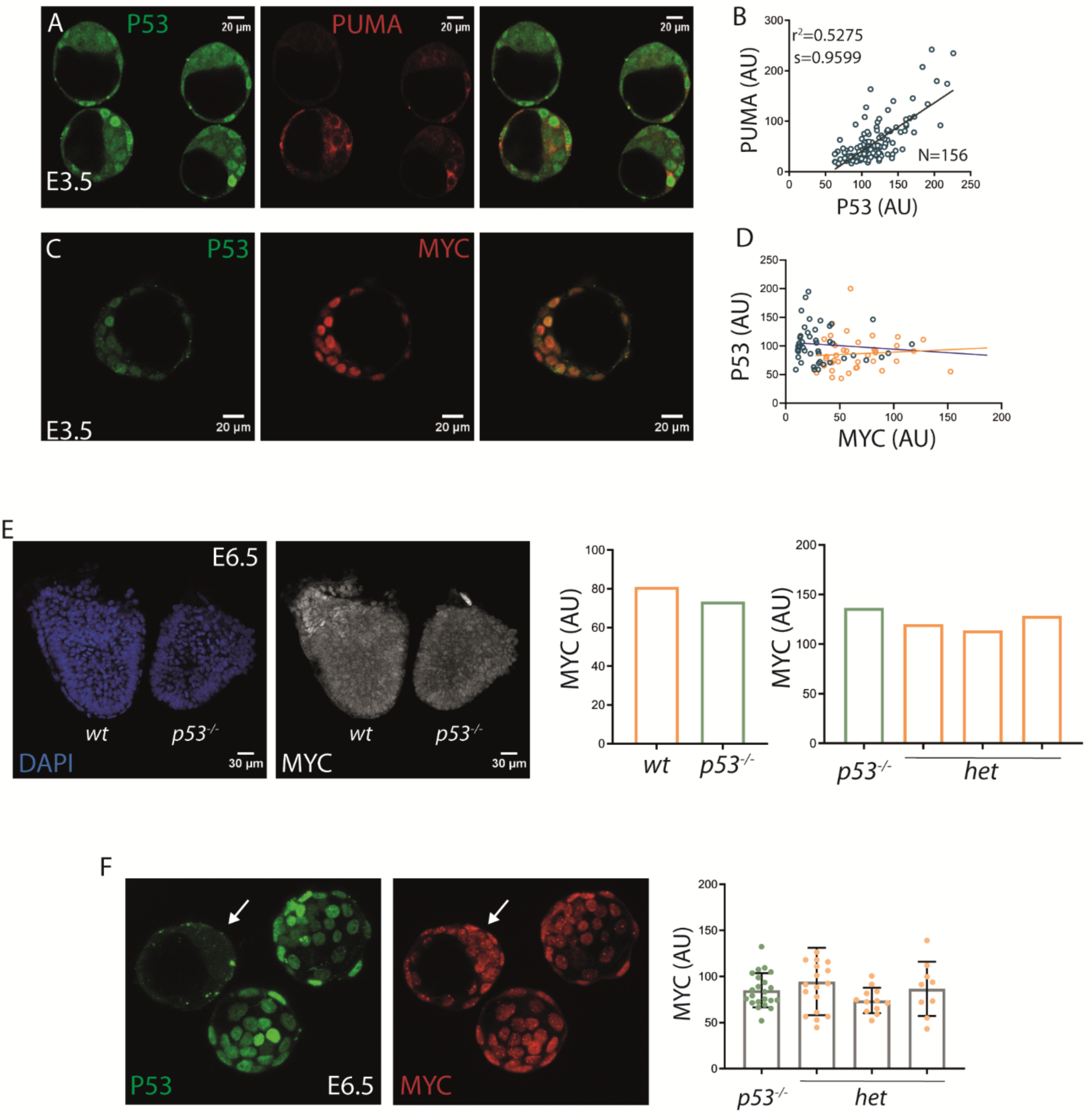
P53, PUMA and MYC expression in the early mouse embryo. **A.** Confocal captures showing P53 and PUMA expression in E3.5 mouse embryos and quantification (**B**). **C.** P53 and MYC expression in E3.5 embryos and quantification (**D**). **E.** MYC expression in *wt* and *p53^-/-^* E6.5 mouse embryos (left) and quantification (right). **F,** Same approach as in **E,** but in E3.5 mouse embryos. White arrow shows a *p53^-/-^* embryo.

**Figure S6.**
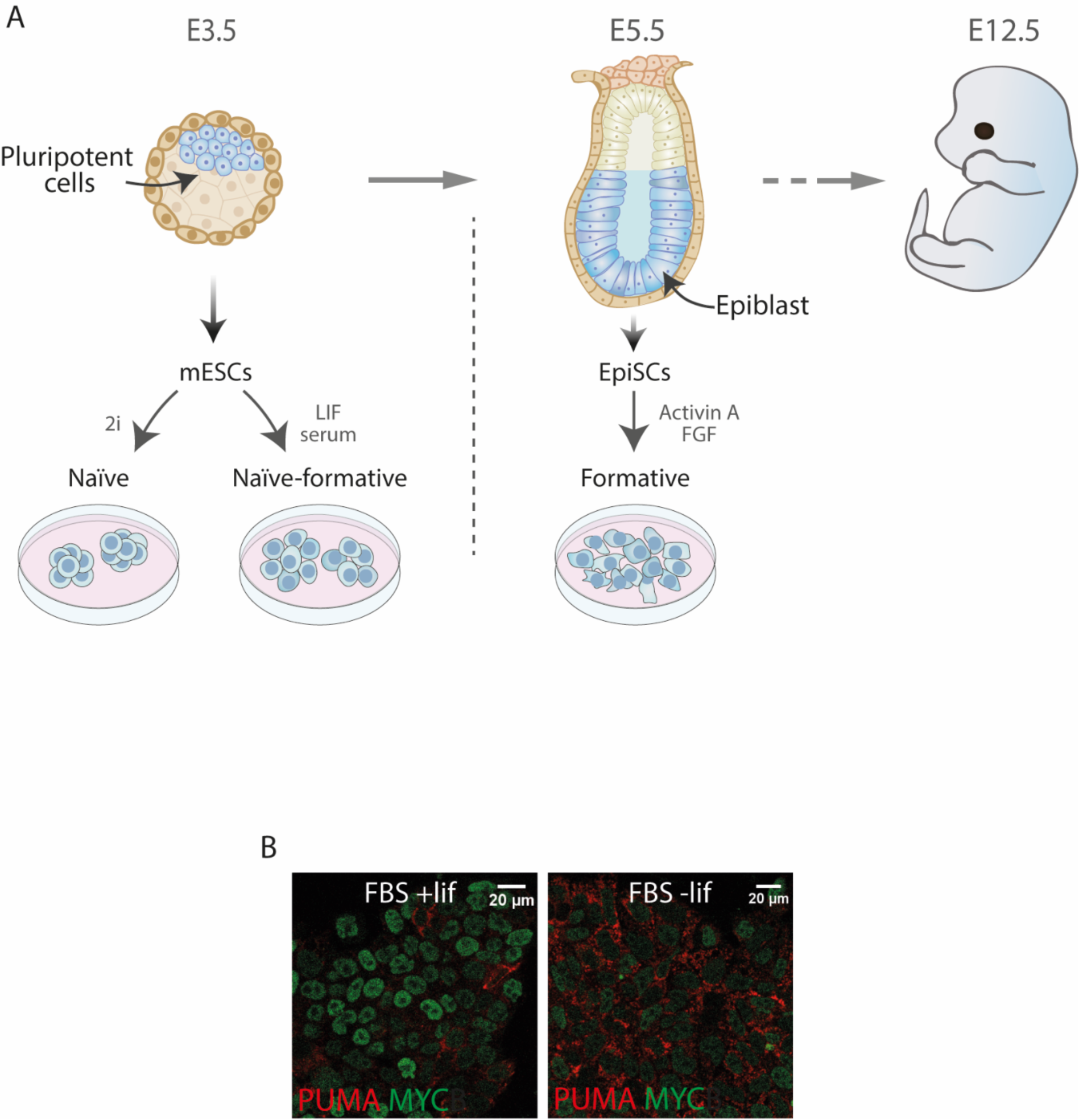
Pluripotent stages *in vivo* and *in vitro.* Pluripotency modulates PUMA and MYC expression. **A.** Scheme representing different pluripotency stages in mouse development, and the *in vitro* models. **B.** PUMA and MYC levels in conventional and differentiating conditions.

**Figure S7.**
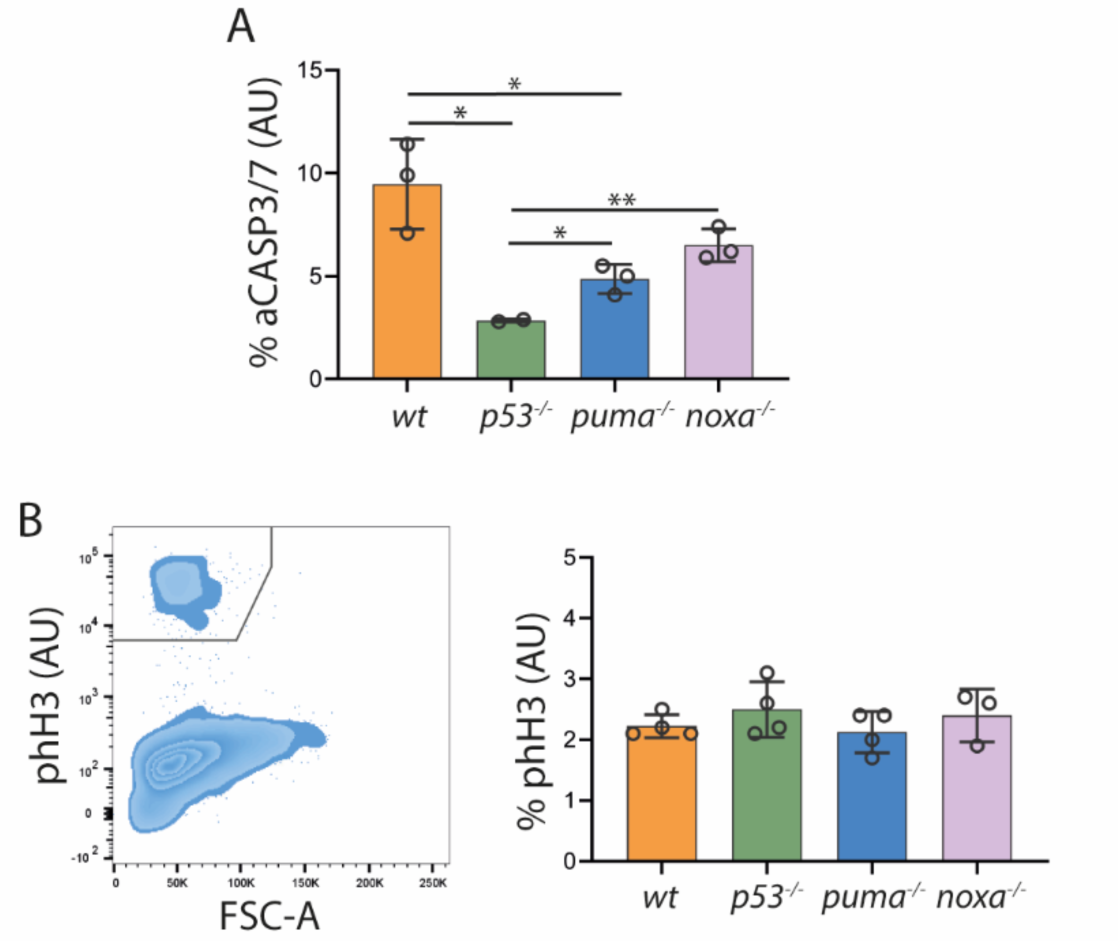
P53, PUMA and NOXA have a role in apoptosis but not in ESC proliferation. **A.** Percentage of active CASP3/7 using the fluorogenic CASP substrate FLICA^TM^. **B.** Contour dot plot showing phH3 positive and negative cells populations (left). Bar graph showing percentage of positive phH3 cells in the indicated ES cell lines. Each dot represents one different clone.

**Figure S8.**
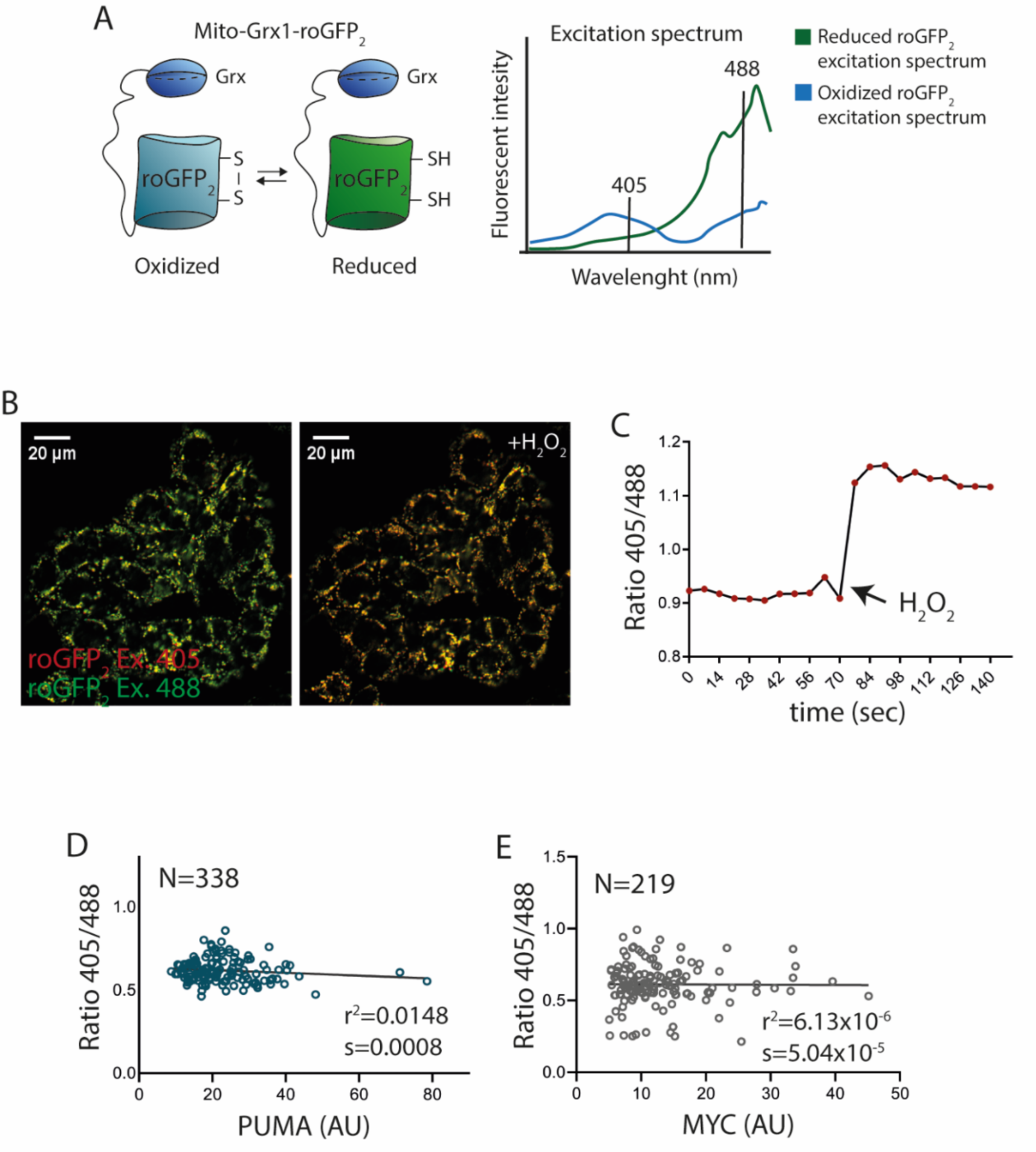
REDOX status in ESCs using the Grx1-roGFP_2_ reporter. **A.** Scheme representing the reporter Grx1-roGFP_2_ in its oxidized and reduced forms (right) and the excitation spectra (left). **B.** roGFP_2_ emission after exciting at 405 and 488nm is represented as a “merge” before and upon H_2_O_2_ treatment and quantification (**C**)**. D, E.** Quantification of REDOX status and PUMA and MYC levels respectively.

**Figure S9.**
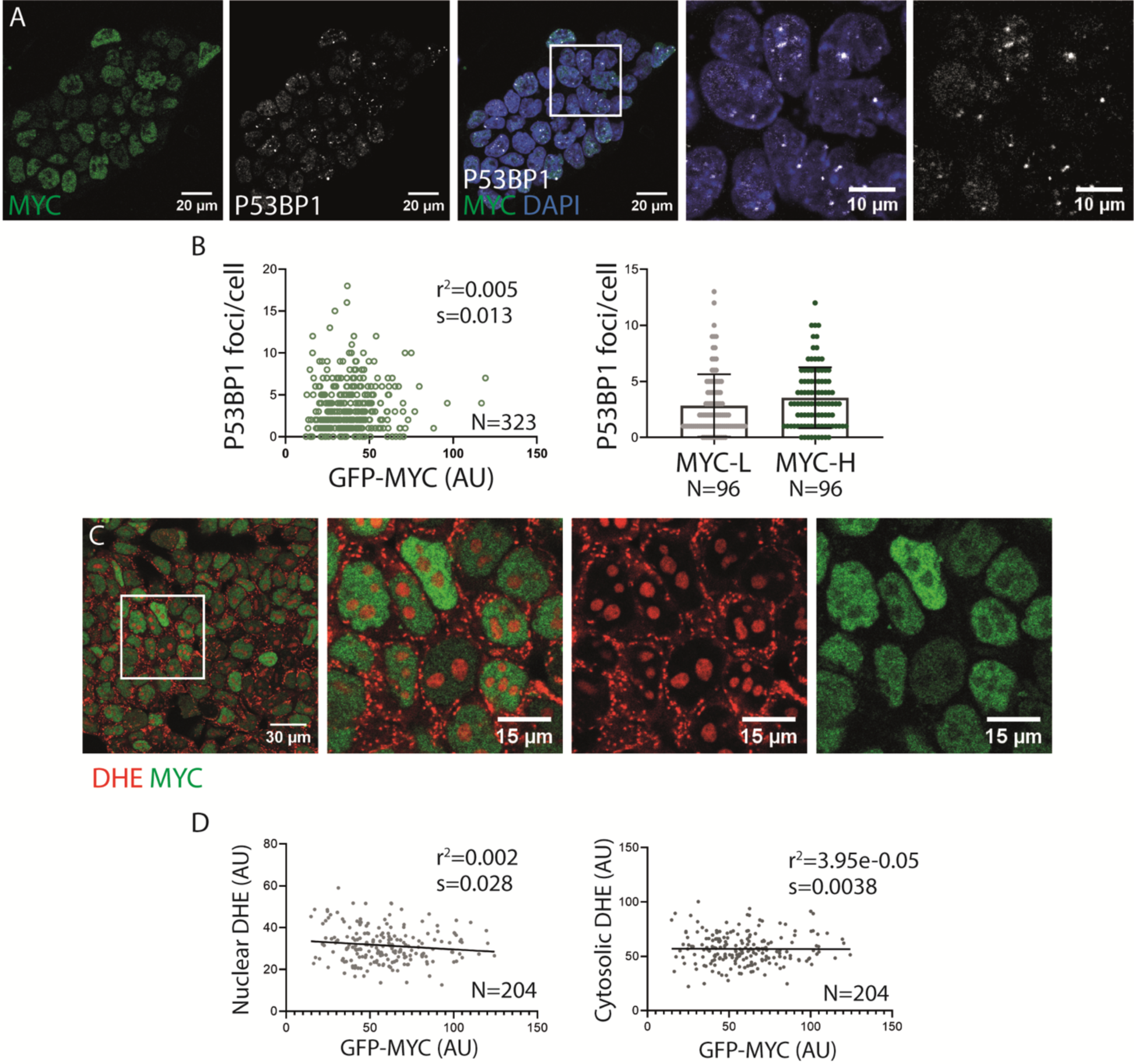
DNA damage and oxidative stress do not correlate with MYC levels. **A.** MYC expression and P53BP1 foci in ES cells and quantification (**B**). **C.** DHE and MYC expression in ESCs and quantification (**D**).

**Figure S10.**
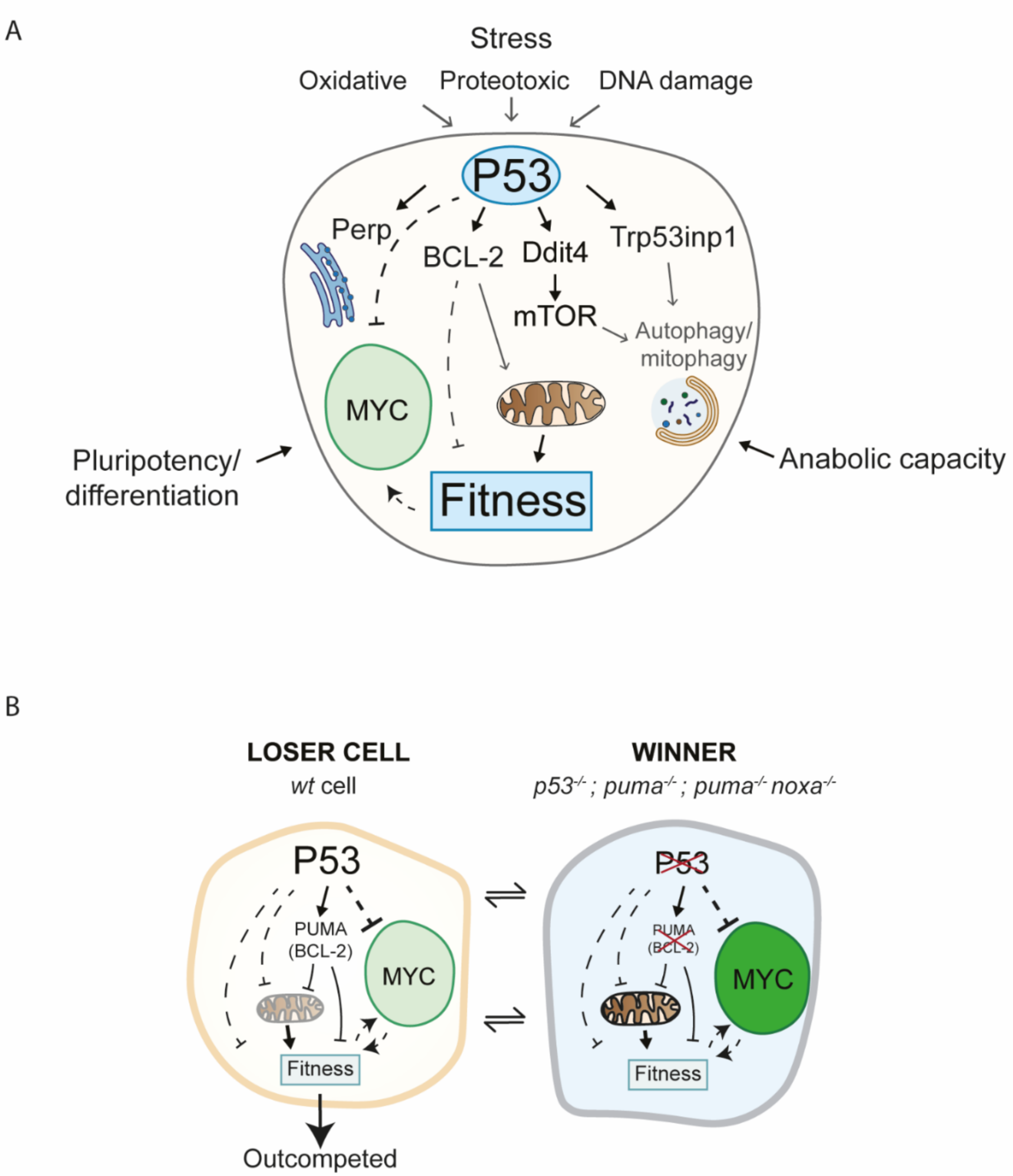
Model. **A.** Stress signature, pluripotent status or anabolic capacity (reported by MYC) have been described as important elements in Pluripotent Cell Competition (Lima *et al*, 2021; Clavería *et al*, 2013; Díaz-Díaz *et al*, 2017). P53 is a well described component in Cell Competition in different models, including pluripotent cells (Dejosez *et al*, 2013; Montero *et al*, 2022) and a sensor of cellular stress. Here, we have identified several candidate genes downstream P53 that can exert a role in the competitive fitness involving different mechanisms as mitochondrial function, autophagy or Ca^2+^ homeostasis. **B.** The absence of P53, puma or the simultaneous deletion of PUMA and NOXA is enough to exert competitive interactions and outcompete *wt* cells. Due to the effect of these proteins regulating the mitochondrial membrane potential and the location of PUMA in the mitochondria we hypothesized that the effect in Fitness could be mediated at least in part through the mitochondria.

## ANNEX I

### 1. CYTOPLASM MACRO

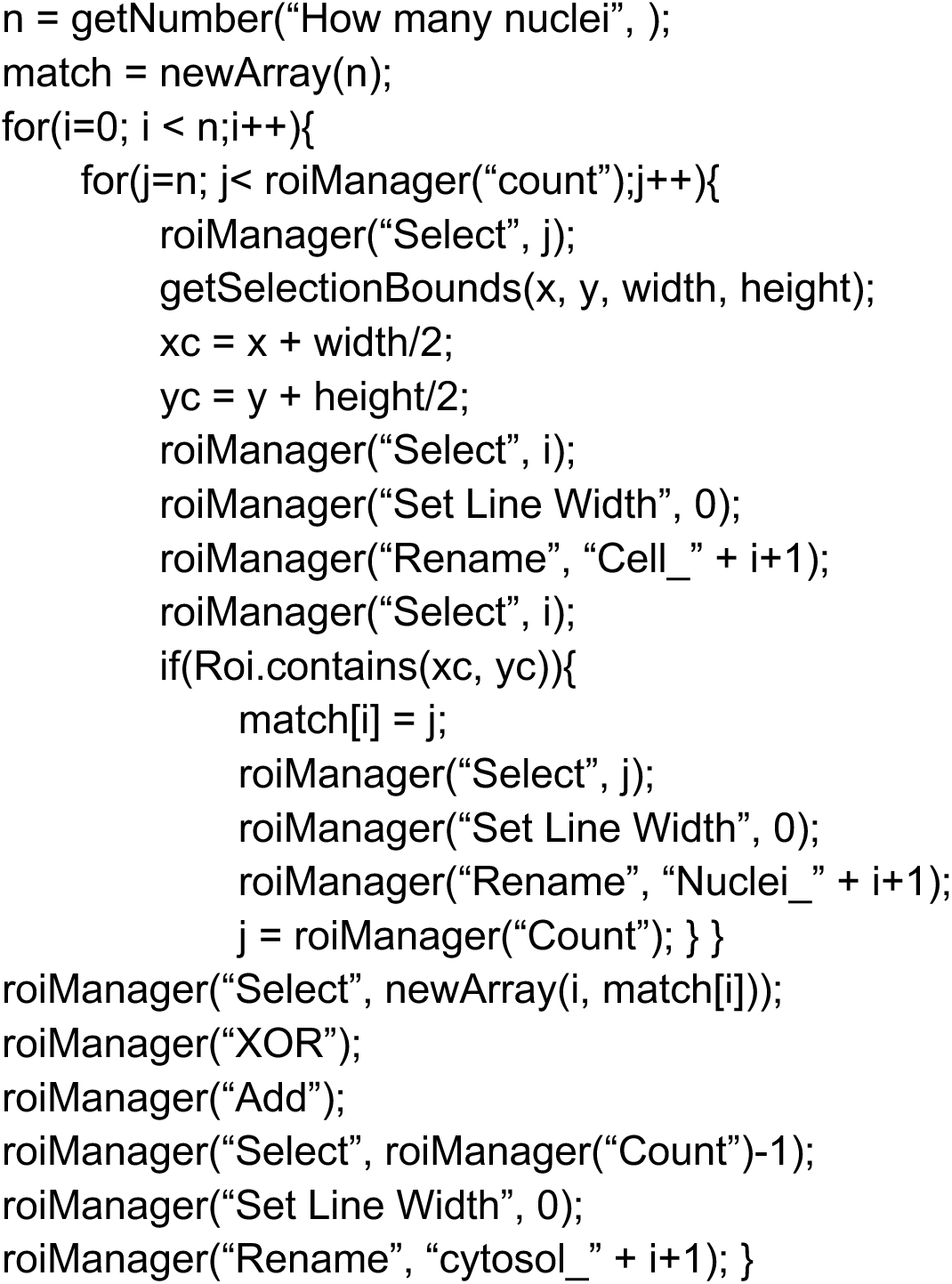

### 2. FOCI NUMBER MACRO

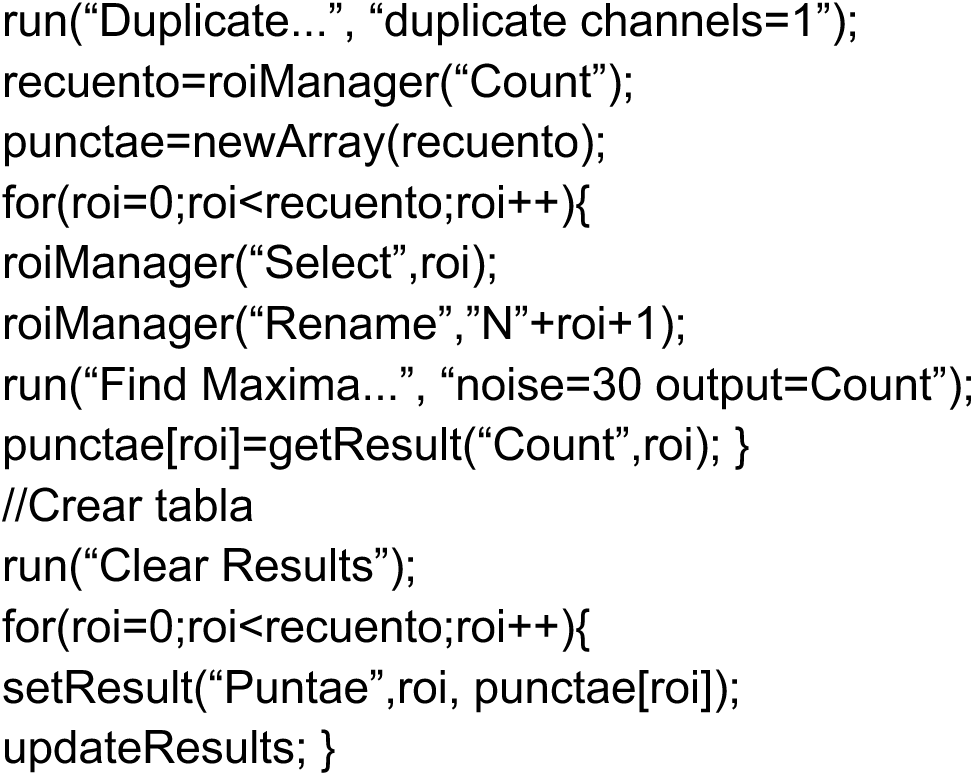

## REFERENCES

Abdelalim EM & Tooyama I (2014) Knockdown of p53 suppresses Nanog expression in embryonic stem cells. Biochem Biophys Res Commun 443: 652–657

Albrecht SC, Barata AG, Großhans J, Teleman AA & Dick TP (2011) In Vivo Mapping of Hydrogen Peroxide and Oxidized Glutathione Reveals Chemical and Regional Specificity of Redox Homeostasis. Cell Metab 14: 819–829

Baumgartner ME, Dinan MP, Langton PF, Kucinski I & Piddini E (2021) Proteotoxic stress is a driver of the loser status and cell competition. Nat Cell Biol 23: 136–146

Bondar T & Medzhitov R (2010) p53-Mediated Hematopoietic Stem and Progenitor Cell Competition. Cell Stem Cell 6: 309–322

Bowling S, Gregorio A Di, Sancho M, Pozzi S, Aarts M, Signore M, Schneider MD, Pedro J, Barbera M, Gil J, et al (2018) P53 and mTOR signalling determine fitness selection through cell competition during early mouse embryonic development. Nat Commun

Bowling S, Lawlor K & Rodr TA (2019) Cell competition : the winners and losers of fitness selection.

Brons IGM, Smithers LE, Trotter MWB, Rugg-Gunn P, Sun B, Chuva de Sousa Lopes SM, Howlett SK, Clarkson A, Ahrlund-Richter L, Pedersen RA, et al (2007) Derivation of pluripotent epiblast stem cells from mammalian embryos. Nature 448: 191–195

Certo M, Del Gaizo Moore V, Nishino M, Wei G, Korsmeyer S, Armstrong SA & Letai A (2006) Mitochondria primed by death signals determine cellular addiction to antiapoptotic BCL-2 family members. Cancer Cell 9: 351–365

Clavería C, Giovinazzo G, Sierra R & Torres M (2013) Myc-driven endogenous cell competition in the early mammalian embryo. Nature 500: 39–44

Clavería C & Torres M (2015) Cell Competition: Mechanisms and Physiological Roles.

Coelho DS, Schwartz S, Merino MM, Hauert B, Topfel B, Tieche C, Rhiner C & Moreno E (2018) Culling Less Fit Neurons Protects against Amyloid-β-Induced Brain Damage and Cognitive and Motor Decline. Cell Rep 25: 3661–3673.e3

Concordet JP & Haeussler M (2018) CRISPOR: Intuitive guide selection for CRISPR/Cas9 genome editing experiments and screens. Nucleic Acids Res 46: W242–W245

Coucouvanis E & Martin GR (1995) Signals for death and survival: a two-step mechanism for cavitation in the vertebrate embryo. Cell 83: 279–287

Deathridge J, Antolović V, Parsons M & Chubb JR (2019) Live imaging of erk signalling dynamics in differentiating mouse embryonic stem cells. Development (Cambridge) 146

Dejosez M (2013) Safeguards for Cell Cooperation in. Science (1979) 341: 1511–4

Dejosez M, Ura H, Brandt VL & Zwaka TP (2013) Safeguards for Cell Cooperation in Mouse Embryogenesis Shown by Genome-Wide Cheater Screen. Science (1979) 341: 1511 LP – 1514

Díaz-Díaz C, Manuel LF De, Jimenez-carretero D, Torres M & Claverıa C (2017) Pluripotency Surveillance by Myc-Driven Competitive Elimination of Differentiating Cells. 585–599

Fernandez-Antoran D, Piedrafita G, Murai K, Ong SH, Herms A, Frezza C & Jones PH (2019) Outcompeting p53-Mutant Cells in the Normal Esophagus by Redox Manipulation. Cell Stem Cell 25: 329–341.e6

Fu X, Wu S, Li B, Xu Y & Liu J (2020) Functions of p53 in pluripotent stem cells. Protein Cell 11: 71–78

Goedhart J & Luijsterburg MS (2020) VolcaNoseR is a web app for creating, exploring, labeling and sharing volcano plots. Sci Rep 10

Gregorio A di, Bowling S & Rodriguez TA (2016) Review Cell Competition and Its Role in the Regulation of Cell Fitness from Development to Cancer. Dev Cell 38: 621–634

Guo G, Pinello L, Han X, Lai S, Shen L, Lin TW, Zou K, Yuan GC & Orkin SH (2016) Serum-Based Culture Conditions Provoke Gene Expression Variability in Mouse Embryonic Stem Cells as Revealed by Single-Cell Analysis. Cell Rep 14: 956–965

Gutscher M, Pauleau A, Marty L, Brach T, Wabnitz GH, Samstag Y, Meyer AJ & Dick TP (2008) Real-time imaging of the intracellular glutathione redox potential. 5: 553–559

Hackett JA & Surani MA (2014) Regulatory principles of pluripotency: from the ground state up. Cell Stem Cell 15: 416–430

Handy DE & Loscalzo J (2012) Redox regulation of mitochondrial function. Antioxid Redox Signal 16: 1323–1367

Hao Q, Chen J, Liao J, Huang Y, Larisch S, Zeng SX, Lu H & Zhou X (2020) p53 induces ARTS to promote mitochondrial apoptosis. bioRxiv: 2020.05.14.096982

Hashimoto M & Sasaki H (2019) Epiblast Formation by TEAD-YAP-Dependent Expression of Pluripotency Factors and Competitive Elimination of Unspecified Cells. Dev Cell 50: 139–154.e5

Heyer BS, Macauley A, Behrendtsen O & Werb Z (2000) Hypersensitivity to DNA damage leads to increased apoptosis during early mouse development. Genes Dev 14: 2072–2084

Huang CY, Bredemeyer AL, Walker LM, Bassing CH & Sleckman BP (2008) Dynamic regulation of c-Myc proto-oncogene expression during lymphocyte development revealed by a GFP-c-Myc knock-in mouse. Eur J Immunol 38: 342–349

Jacks T, Remington L, Williams B 0, Schmitt EM, Halachmit S, Bronson RT & Weinberg RA (1994) Tumor spectrum analysis in p53-mutant mice

Jain AK, Allton K, Iacovino M, Mahen E, Milczarek RJ, Zwaka TP, Kyba M & Barton MC (2012) p53 regulates cell cycle and microRNAs to promote differentiation of human embryonic stem cells. PLoS Biol 10: e1001268

Jain AK & Barton MC (2018) p53: emerging roles in stem cells, development and beyond. Development 145

Jaiswal SK, Oh JJ & DePamphilis ML (2020) Cell cycle arrest and apoptosis are not dependent on p53 prior to p53-dependent embryonic stem cell differentiation. Stem Cells 38: 1091–1106

Kastenhuber ER & Lowe SW (2017) Putting p53 in Context. Cell 170: 1062–1078

Kim J, Yu L, Chen W, Xu Y, Wu M, Todorova D, Tang Q, Feng B, Jiang L, He J, et al (2019) Wild-Type p53 Promotes Cancer Metabolic Switch by Inducing PUMA-Dependent Suppression of Oxidative Phosphorylation. Cancer Cell 35: 191–203.e8

Kinoshita M, Barber M, Mansfield W, Cui Y, Spindlow D, Stirparo GG, Dietmann S, Nichols J & Smith A (2021) Capture of Mouse and Human Stem Cells with Features of Formative Pluripotency. Cell Stem Cell 28: 453–471.e8

Kon S & Fujita Y (2021) Cell competition-induced apical elimination of transformed cells, EDAC, orchestrates the cellular homeostasis. Dev Biol 476: 112–116 doi:10.1016/j.ydbio.2021.03.015 [PREPRINT]

Kucinski I, Dinan M, Kolahgar G & Piddini E (2017) Chronic activation of JNK JAK/STAT and oxidative stress signalling causes the loser cell status. Nat Commun 8

Levayer R & Moreno E (2013) Mechanisms of cell competition: Themes and variations. J Cell Biol 200: 689–698

Lima A, Lubatti G, Burgstaller J, Hu D, Green AP, Di Gregorio A, Zawadzki T, Pernaute B, Mahammadov E, Perez-Montero S, et al (2021) Cell competition acts as a purifying selection to eliminate cells with mitochondrial defects during early mouse development. Nat Metab 3: 1091–1108

Lin T, Chao C, Saito S, Mazur SJ, Murphy ME, Appella E & Xu Y (2005) p53 induces differentiation of mouse embryonic stem cells by suppressing Nanog expression. Nat Cell Biol 7: 165–171

Manova K, Tomihara-Newberger C, Wang S, Godelman A, Kalantry S, Witty-Blease K, De Leon V, Chen WS, Lacy E & Bachvarova RF (1998) Apoptosis in mouse embryos: Elevated levels in pre gastrulae and in the distal anterior region of gastrulae of normal and mutant mice. Developmental Dynamics 213: 293–308

Marusyk A, Porter CC, Zaberezhnyy V & DeGregori J (2010) Irradiation Selects for p53-Deficient Hematopoietic Progenitors. PLoS Biol 8: e1000324

McDonnell SJ, Spiller DG, White MRH, Prior IA & Paraoan L (2019) ER stress-linked autophagy stabilizes apoptosis effector PERP and triggers its co-localization with SERCA2b at ER– plasma membrane junctions. Cell Death Discov 5

Merino MM, Rhiner C, Lopez-gay JM, Buechel D, Hauert B & Moreno E (2015) Article Elimination of Unfit Cells Maintains Tissue Health and Prolongs Lifespan. Cell 160: 461–476

Montero SP, Bowling S, Pérez-Carrasco R & Rodriguez TA (2022) Levels of p53 expression determine the competitive ability of embryonic stem cells during the onset of differentiation. bioRxiv: 2022.02.28.482311

Morata G & Calleja M (2020) Cell competition and tumorigenesis in the imaginal discs of Drosophila. Semin Cancer Biol 63: 19–26 doi:10.1016/j.semcancer.2019.06.010 [PREPRINT]

Nichols J & Smith A (2009) Naive and primed pluripotent states. Cell Stem Cell 4: 487–492

Østergaard HHAHFG; WJR (2001) Shedding light on disulfide bond formation: engineering a redox switch in green fluorescent protein. EMBO JOURNAL 20: 5853–5862

Pernaute B, Pérez-Montero S, Sánchez Nieto JM, Di Gregorio A, Lima A, Lawlor K, Bowling S, Liccardi G, Tomás A, Meier P, et al (2022) DRP1 levels determine the apoptotic threshold during embryonic differentiation through a mitophagy-dependent mechanism. Dev Cell 57: 1316–1330.e7

Pernaute B, Spruce T, Smith KM, Sánchez-Nieto JM, Manzanares M, Cobb B & Rodríguez TA (2014) MicroRNAs control the apoptotic threshold in primed pluripotent stem cells through regulation of BIM. Genes Dev 28: 1873–1878

Posfai E, Tam OH & Rossant J (2014) Chapter One - Mechanisms of Pluripotency In Vivo and In Vitro. In Stem Cells in Development and Disease, Rendl MBT-CT in DB (ed) pp 1–37. Academic Press

Raudvere U, Kolberg L, Kuzmin I, Arak T, Adler P, Peterson H & Vilo J (2019) G:Profiler: A web server for functional enrichment analysis and conversions of gene lists (2019 update). Nucleic Acids Res 47: W191–W198

Saadi H, Seillier M & Carrier A (2015) The stress protein TP53INP1 plays a tumor suppressive role by regulating metabolic homeostasis. Biochimie 118: 44–50

Sancho M, Di-Gregorio A, George N, Pozzi S, Sánchez JM, Pernaute B & Rodríguez TA (2013) Competitive interactions eliminate unfit embryonic stem cells at the onset of differentiation. Dev Cell 26: 19–30

Sato N, Meijer L, Skaltsounis L, Greengard P & Brivanlou AH (2004) Maintenance of pluripotency in human and mouse embryonic stem cells through activation of Wnt signaling by a pharmacological GSK-3-specific inhibitor. Nat Med 10: 55–63

Schindelin J, Arganda-Carreras I, Frise E, Kaynig V, Longair M, Pietzsch T, Preibisch S, Rueden C, Saalfeld S, Schmid B, et al (2012) Fiji: An open-source platform for biological-image analysis. Nat Methods 9: 676–682 doi:10.1038/nmeth.2019 [PREPRINT]

Siddiqui WA, Ahad A & Ahsan H (2015) The mystery of BCL2 family: Bcl-2 proteins and apoptosis: an update. Arch Toxicol 89: 289–317

Singla S, Iwamoto-Stohl LK, Zhu M & Zernicka-Goetz M (2020) Autophagy-mediated apoptosis eliminates aneuploid cells in a mouse model of chromosome mosaicism. Nat Commun 11

Sperber H, Mathieu J, Wang Y, Ferreccio A, Hesson J, Xu Z, Fischer KA, Devi A, Detraux D, Gu H, et al (2015) The metabolome regulates the epigenetic landscape during naive-to-primed human embryonic stem cell transition. Nat Cell Biol 17: 1523–1535

Tesar PJ, Chenoweth JG, Brook FA, Davies TJ, Evans EP, Mack DL, Gardner RL & McKay RDG (2007) New cell lines from mouse epiblast share defining features with human embryonic stem cells. Nature 448: 196–199

Tirado-Hurtado I, Fajardo W & Pinto JA (2018) DNA damage inducible transcript 4 gene: The switch of the metabolism as potential target in cancer. Front Oncol 8

Wagstaff L, Goschorska M, Kozyrska K, Duclos G, Kucinski I, Chessel A, Neil LH, Bradshaw CR, Allen GE, Rawlins EL, et al (2016) Mechanical cell competition kills cells via induction of lethal p53 levels.

Wagstaff L, Kolahgar G & Piddini E (2013) Competitive cell interactions in cancer : a cellular tug of war. Trends Cell Biol 23: 160–167

Ying QL, Wray J, Nichols J, Batlle-Morera L, Doble B, Woodgett J, Cohen P & Smith A (2008) The ground state of embryonic stem cell self-renewal. Nature 453: 519–523

Yu J & Zhang L (2008) PUMA, a potent killer with or without p53. Oncogene 27: 71–83

Zhang G, Xiea Y, Zhou Y, Xiang C, Chen L, Zhang C, Hou X, Chen J, Zong H & Liu G (2017) P53 pathway is involved in cell competition during mouse embryogenesis. Proc Natl Acad Sci U S A 114: 498–503

Zhang Z-N, Chung S-K, Xu Z & Xu Y (2014) Oct4 maintains the pluripotency of human embryonic stem cells by inactivating p53 through Sirt1-mediated deacetylation. Stem Cells 32: 157–165

